# Circular-Arc Modal Migration for Ultrasound Imaging—Part I: Finite-Arc Spectral Theory and Stolt–Pullback Factorization

**DOI:** 10.64898/2026.11.11.765911

**Authors:** Zihan Huang, Fan Shi, Dawei Wu, Chuan Chen

## Abstract

Circular-arc ultrasound imaging requires a spectral construction that accommodates radius-dependent propagation and the spatially varying phase of offset-source transmission. This first of two articles develops Circular-Arc Modal Migration (CAMM), a direct Stolt-type imaging framework in angular modal coordinates. Starting from a first-Born acoustic model, angular decomposition and local radial Stolt mappings convert channel spectra into fields for coherent image synthesis. The offset transmit phase is represented by a phase-screen factorization through reusable spectral factors and spatially weighted fields. A scalar-pullback approximation simplifies this correction to reading one reference image at a geometrically fitted radial coordinate. Both realizations share the same modal construction. Error controls address local radial synthesis, transmit-path expansion, scalar approximation, and phase-screen truncation, with direct integration verifying the separated numerical evaluation. Same-RF point-target comparisons show that scalar pullback closely preserves the principal peak positions and widths of phase-screen imaging in the compact semicircular configuration, while phase screens better control surrounding responses in the shallow convex examples. The retained modal spectrum, frequency band, image radius, and source displacement guide this accuracy–computation choice. CAMM thereby extends direct spectral image formation to circular arcs and supplies the theoretical and numerical basis for the image-quality and performance studies in Part II.

## I. Introduction

**C**IRCULAR-ARC ultrasound arrays combine a broad angular field with a compact probe footprint. Conventional convex probes provide sector imaging [1], while divided-ring and forward-viewing ring arrays support volumetric and interventional imaging [2]–[4]. Forward-looking catheter arrays further illustrate the need to form images through restricted probe footprints [5], [6]. For outward-looking pulse-echo acquisitions, synthetic-aperture processing [7] and coherent compounding of plane or diverging waves [8], [9] combine information from multiple illuminations. More emissions increase acquisition duration and exposure to inter-event motion [10]. A useful reconstruction must therefore exploit the acquired views across the sector, concentrating reflector responses while limiting signal outside the target region. The broad angular coverage makes response variation with depth and angle important. This motivates examining how the reconstruction distributes signal around reflectors as their positions change.

Delay-and-sum (DAS) forms each image point by aligning and summing channel samples according to its transmit and receive paths. Its image quality depends on the receive aperture and propagation parameters [11], as well as the transmit-field representation used in synthetic-aperture reconstruction [12]. In the compact semicircular-array examples studied here, identifiable DAS peaks coexist with extended arc-shaped off-target responses. Peak localization and the surrounding signal therefore describe complementary aspects of the spatial response. Related convex-array studies show that a transmit arrangement producing a narrow main lobe can also produce strong secondary responses for oblique targets and reduce cyst contrast [13]; diverging-wave source placement likewise changes the balance between resolution and sidelobes [14]. A reconstruction intended for a wide sector should thus be evaluated through both the localized response and the surrounding signal.

Existing beamforming approaches address different aspects of the image response. Generalized coherence and phase-coherence weighting modify channel contributions according to their agreement [15], [16], while short-lag coherence imaging forms a coherence-based image statistic [17]. Minimum-variance beamforming adapts the channel weights [18], [19], and delay-multiply-and-sum changes the channel combination [20]. These approaches modify the weighting, combination, or statistical interpretation of the received signals.

Another line of work formulates propagation-based image formation in the spectral domain. Stolt migration maps temporal frequency to an image wavenumber through the propagation dispersion relation [21]. Ultrasound implementations include Fourier imaging with limited-diffraction beams [22], frequency-domain synthetic-aperture focusing [23], multilayer omega–*k* migration [24], and plane-wave Stolt imaging [25].

Fourier-domain beamforming can also reduce the sampling burden associated with time-domain processing [26]. These representative direct constructions use linear or planar acquisition coordinates. For a linear aperture, a Fourier component is indexed by a transverse spatial wavenumber. On a circular arc, the corresponding coordinate is angle: a pattern of fixed angular order spans a longer physical arc at a larger radius. Its tangential wavenumber therefore decreases with radius, and its radial propagation relation changes accordingly. The frequency-to-radial-wavenumber relation is consequently tied to the reconstruction radius as well as the angular order. Adapting direct spectral migration to circular arcs therefore requires a radial mapping that follows this variation while retaining the propagation phase for image synthesis.

Angular Fourier modes provide coordinates adapted to the receive arc. Circular-harmonic beamforming uses this representation to address directivity and sidelobe constraints [27]. For pulse-echo imaging, Ali and Dahl’s polar angular-spectrum method (ASM) extends Fourier beamforming to curvilinear arrays by radially extrapolating transmit and receive wavefields and combining them through zero-lag correlation [1]. This provides a wavefield-based alternative to virtual-source DAS. A complementary Stolt-type construction would map the measured channel spectra directly to radial image coordinates. For a circular arc, this mapping must account for the radius-dependent propagation relation. The resulting question is how to construct that direct spectral image while retaining the phase associated with each transmit event.

Offset-source transmission adds a position-dependent phase to this construction. A source at the curvature center has the same transmit distance to all points on a given radius. Moving the source makes that distance depend on both radius and angle. Relative to centered transmission, the additional path produces a phase correction that varies across the image and between transmit events. Retaining this correction aligns the reconstructed complex fields for coherent compounding. Its position dependence also has to be represented within the radial spectral synthesis, linking the circular-arc remapping problem to the treatment of offset illumination.

We propose Circular-Arc Modal Migration (CAMM), a direct spectral imaging method that combines angular modal analysis, local radial Stolt remapping, and offset-phase compensation. Local radial intervals accommodate the changing propagation relation, and overlapping interval images form the reconstruction. Within each interval, a separable phase-screen approximation represents the position-dependent transmit correction through radial field synthesis and spatially varying complex weights. A simpler scalar-pullback implementation evaluates a reference field at a corrected radial coordinate. The phase-screen implementation retains residual mode-and frequency-dependent effects beyond this coordinate correction by synthesizing additional fields. Both implementations are compared with DAS on the same RF data in two outward-looking circular-arc configurations: a compact semicircular array and a conventional convex array. The former has a 0.75-mm radius and nominal 180^*°*^ angular aperture; the latter has a mm radius and 70^*°*^ aperture. These contrasting scales test how much offset-phase detail is needed to preserve the central and surrounding response. Supplementary engineering examples relate the implementation choice to source displacement, image radius, retained frequency band, and angular spectrum. This two-part study develops and evaluates the CAMM framework. Part I establishes the imaging formulation and verifies its numerical evaluation through component tests, comparison with direct coupled integration, and point-target reconstruction. Part II examines image quality and computational performance through extended simulations and convex-array measurements, including the spatial distribution of point responses and separation of low-echo targets from their backgrounds.

Section II introduces the acquisition geometry and imaging formulation. Section III develops CAMM, and Section IV evaluates its approximations and point responses. Sections V and VI discuss the findings and conclude the paper. The appen-dices provide supporting derivations; supplementary material presents further numerical controls and engineering selection examples.

## II. Geometry and Imaging Formulation

The proposed image construction starts from the transmit and receive paths. The forward model relates these paths, their propagation amplitudes, and the scattering contrast to measured channel data. Its phase then supplies the reference and offset correction used by modal migration. Table I lists the principal symbols; additional implementation quantities are defined where used in the supplement.

**TABLE I.**
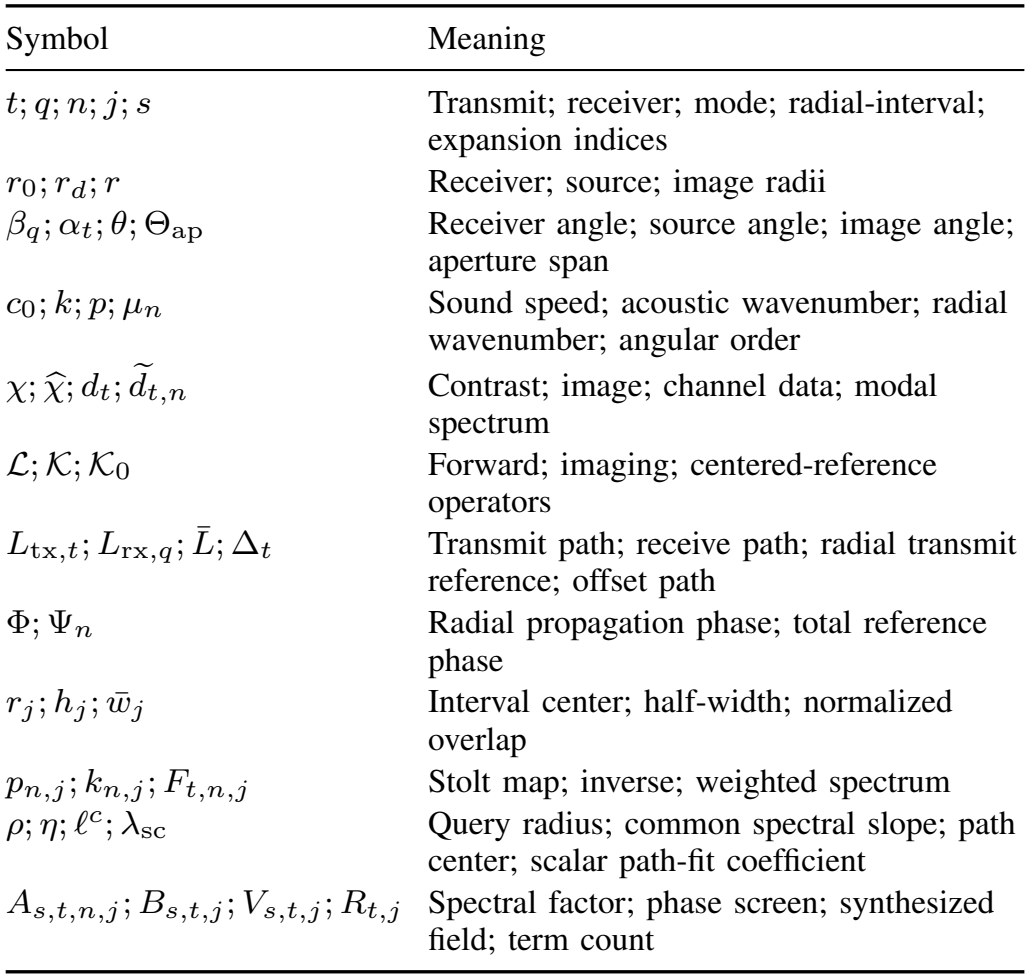
Core Notation.

| Symbol | Meaning |
|---|---|
| t; q; n; j; s | Transmit; receiver; mode; radial-interval; expansion indices |
| r<sub>0</sub>; r<sub>d</sub>; r | Receiver; source; image radii |
| β<sub>q</sub>; α<sub>t</sub>; θ; Θ<sub>ap</sub> | Receiver angle; source angle; image angle; aperture span |
| c<sub>0</sub>; k; p; μ<sub>n</sub> | Sound speed; acoustic wavenumber; radial wavenumber; angular order |
| $\chi; \widehat{\chi}; d_t; \widetilde{d}_{t,n}$ | Contrast; image; channel data; modal spectrum |
| $\mathcal{L}; \mathcal{K}; \mathcal{K}_0$ | Forward; imaging; centered-reference operators |
| $L_{\mathrm{tx},t}; L_{\mathrm{rx},q}; \bar{L}; \Delta_t$ | Transmit path; receive path; radial transmit reference; offset path |
| Φ; Ψ<sub>n</sub> | Radial propagation phase; total reference phase |
| $r_j; h_j; \bar{w}_j$ | Interval center; half-width; normalized overlap |
| p<sub>n,j</sub>; k<sub>n,j</sub>; F<sub>t,n,j</sub> | Stolt map; inverse; weighted spectrum |
| ρ; η; ℓ<sup>c</sup>; λ<sub>sc</sub> | Query radius; common spectral slope; path center; scalar path-fit coefficient |
| A<sub>s,t,n,j</sub>; B<sub>s,t,j</sub>; V<sub>s,t,j</sub>; R<sub>t,j</sub> | Spectral factor; phase screen; synthesized field; term count |

### A. Circular-Arc Acquisition

Consider an outward-looking array in a homogeneous background of sound speed *c*_0_. With the curvature center as origin and the symmetry axis at angle zero, the target, receiver, and virtual-source positions are

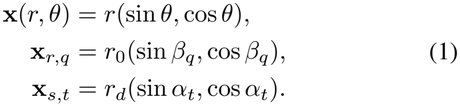

The indices *q* and *t* identify receivers and transmit events. The receive arc spans Θ_ap_; the reconstruction sector Ω lies outside both radii, *r >* max(*r*_0_, *r*_*d*_), and may use a smaller angular field. Figure 1 defines the geometry. The propagation distances are

**Fig. 1.**
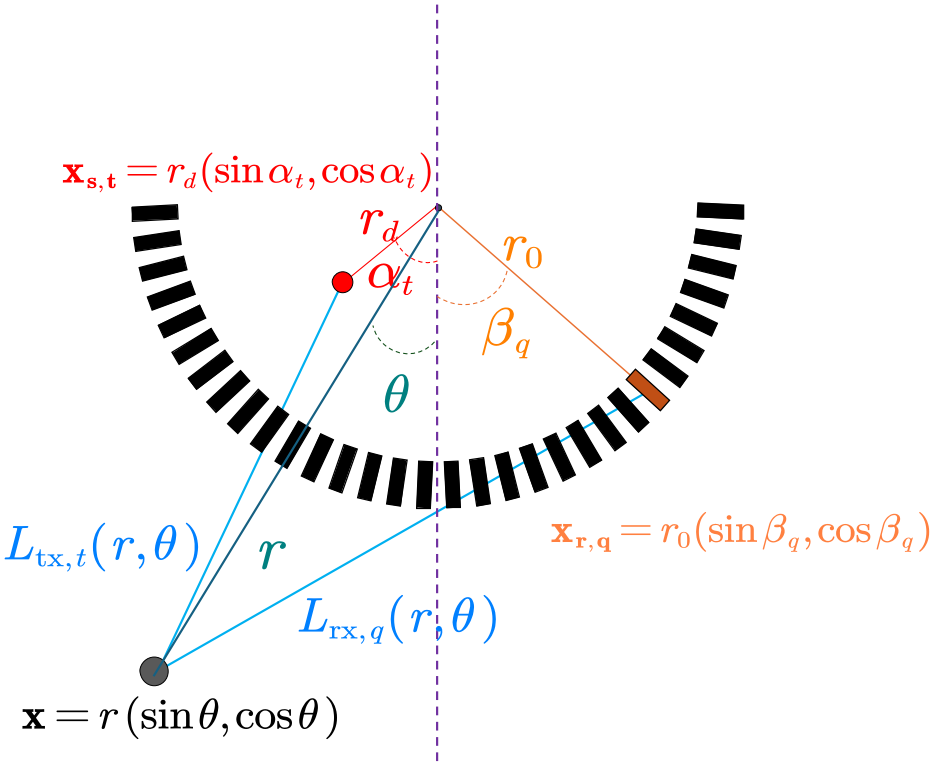
Circular-arc acquisition geometry. The offset source, receiver, and target define the transmit and receive paths. The semicircular sketch illustrates the radii and angles also used for a finite convex arc.

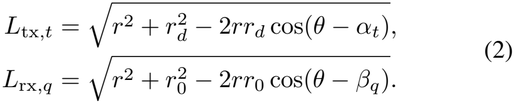

For convex-array displays, depth from the apex is *z* = *r* cos *θ* −*r*_0_; the radial coordinate *r* remains measured from the curvature center throughout the derivation.

### B. From the Wave Equation to the Forward Operator

With these source and receiver positions fixed, let *c*(**x**) be the local sound speed and 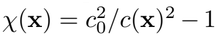 the scattering contrast in a constant-density acoustic model. The pressure *u*_*t*_ generated by a prescribed source *f*_*t*_ at time *τ* satisfies

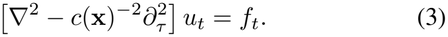

For a temporal signal *v*(*τ*), we use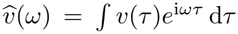. Spectra below are parameterized by *k* = *ω/c*_0_ *>* 0, where *ω* is angular frequency. The outgoing two-dimensional Green function 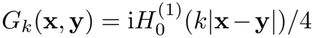 describes propagation from source point **y** to observation point 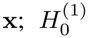 is the Hankel function of the first kind. The first-Born approximation replaces the field illuminating a scatterer by the incident field *S*_*t*_(*k*)*G*_*k*_(**x, x**_*s,t*_), with source spectrum *S*_*t*_. Including the transmit–receive system response *H*_*tq*_ gives the forward operator

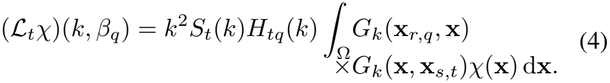

To construct modal migration, we expose the transmit and receive phases through the high-frequency Green-function approximation, following the travel-phase description of migration [28], [29]. Collecting the source spectrum, system response, geometrical spreading, and frequency prefactors into *a*_*t*_ gives the phase-form forward operator Q_*t*_:

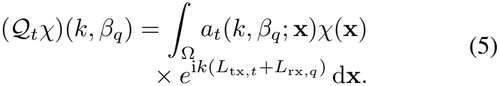

The integral is over scatterer positions in Ω; the receiver position remains indexed by *q*. This separates the weak-scattering assumption in (4) from the high-frequency phase approximation in (5).

### C. The Imaging-Operator Choice

The forward model predicts channel data; image formation estimates the object from those data. Let *d*_*t*_(*k, β*_*q*_) denote the measured spectrum for transmit event *t* = 1, …, *N*_tx_, distinct from time *τ*. Write **d** = (*d*_*t*_)_*t*_ and ℒ*χ* = (_*t*_*χ*)_*t*_ for the data and prediction collections. With measurement error ***ϵ***, the linear data model is

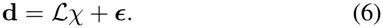

Reconstruction seeks *χ* from **d** with ℒ specified by the acquisition model. Figure 2 compares three representative routes.

**Fig. 2.**
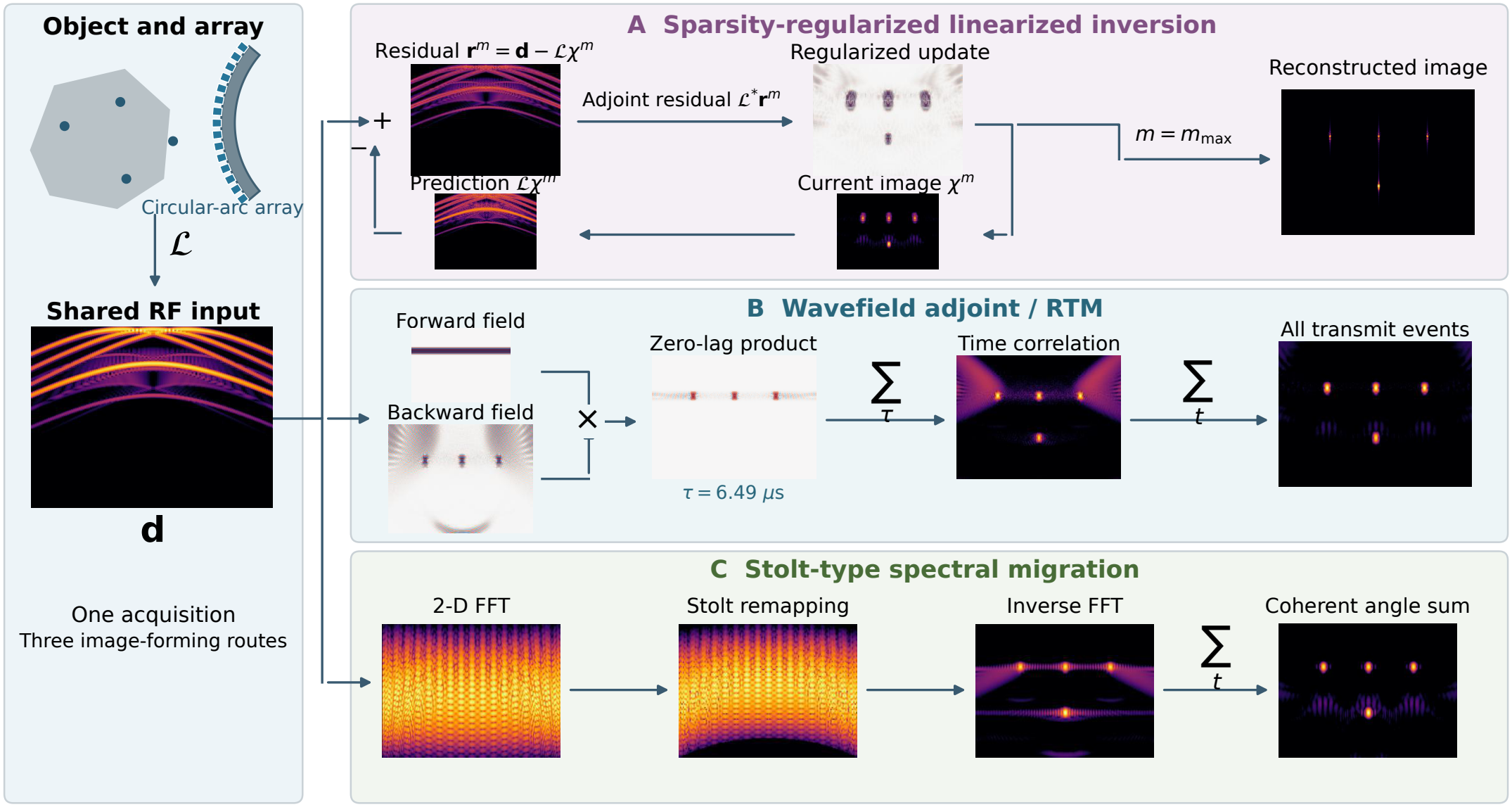
Three imaging routes from channel data: (A) sparsity-regularized linearized inversion; (B) wavefield adjoint/RTM; (C) Stolt-type spectral migration. The intermediate images illustrate the operations. Route C shows the general spectral strategy; the CAMM construction is summarized in Fig. 3.

Route A fits predicted data to measurements, as in least-squares migration [30]–[32] and regularized beamforming [33]. The illustrated choice uses a nonnegative sparsity parameter *λ*_inv_:

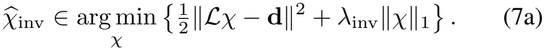

Route B applies the adjoint, implemented through wavefield propagation and correlation as in RTM [34]:

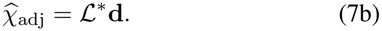

Route C constructs a Stolt-type spectral migration operator and applies it directly:

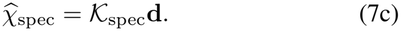

The sparsity-regularized solution in (7a) generally depends nonlinearly on **d** through the regularization. Routes B and C apply fixed linear image maps, written as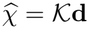. In B, the adjoint returns the data to image space through propagation and correlation; in C, Fourier analysis, spectral remapping, and inverse synthesis form the image directly. Here direct denotes noniterative image formation. Route C follows the established Stolt strategy used in synthetic-aperture and plane-wave ultrasound imaging [21], [23], [25]. CAMM develops this route for circular arcs by combining angular modes with a transmit-dependent modal integral. The next section constructs that integral and two ways to evaluate its offset phase.

## III. Circular-Arc Modal Migration

CAMM first resolves the receive data into angular patterns and remaps their frequency spectra for radial synthesis. The offset transmit path then determines where to sample the synthesized complex fields and how to combine them. Phase-screen evaluation retains the residual coupling through several weighted fields; scalar pullback approximates the correction by sampling one reference image at a shifted radius. Both produce single-transmit images for coherent compounding. Figure 3 provides a reading guide: its operations lead to the equations and derivations developed below.

**Fig. 3.**
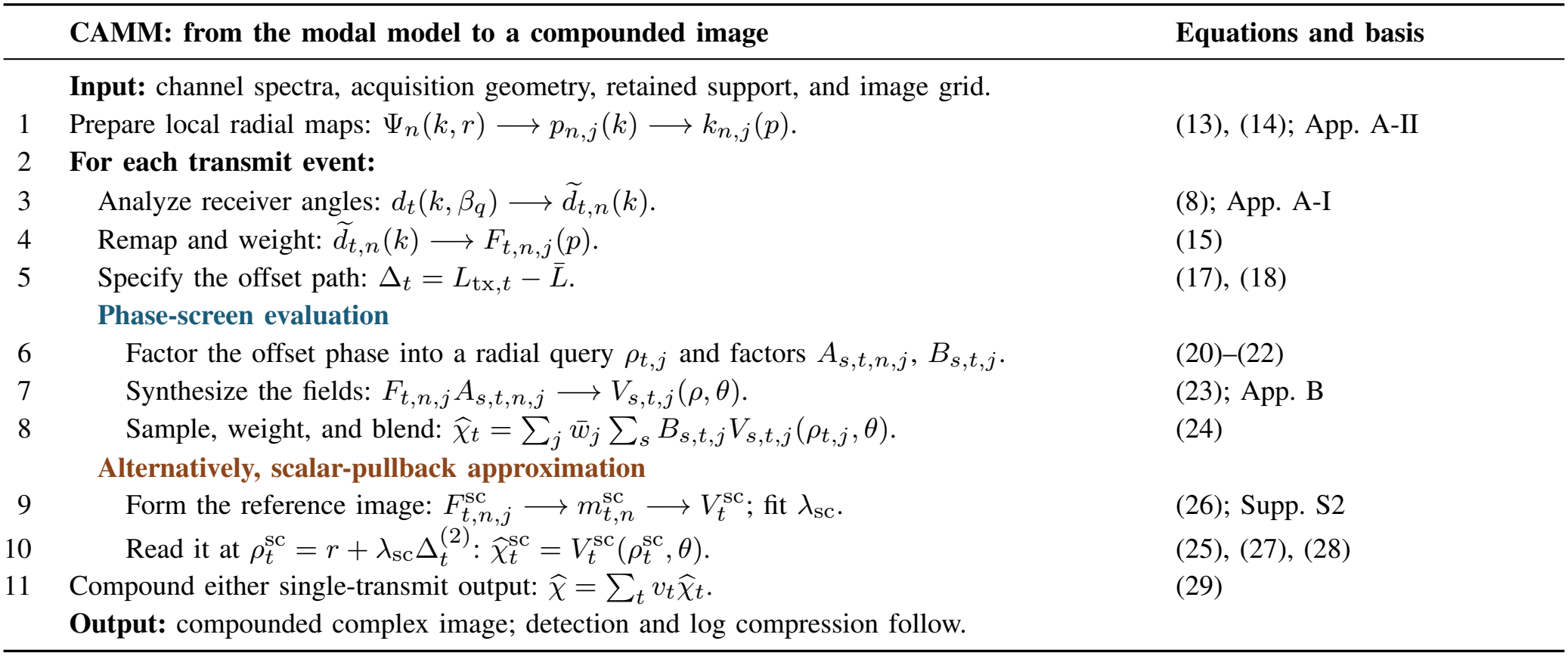
Reconstruction guide with data and symbol transitions. Blue and brown headings identify phase-screen and scalar-pullback evaluation. The query factorization is exact; finite-term screens and scalar path/displacement introduce their respective approximations. The scalar branch forms its own reference field and mode-dependent overlap, as specified in Supplementary Section S2. Both branches provide complex images for coherent compounding.

### A. Angular Modes and Radial Propagation

The receive distance in (2) depends on the angular difference *β* −*θ*, which motivates an angular Fourier representation. We choose the computational period *I*_per_ = [− Θ_ap_*/*2, Θ_ap_*/*2) and orders *µ*_*n*_ = 2*πn/*Θ_ap_; a semicircular array has *µ*_*n*_ = 2*n*. The measured data are analyzed in these angular patterns:

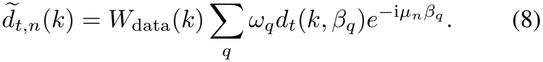

Here *W*_data_ is a common frequency-band multiplier. Channel-dependent calibration, when available, precedes this analysis. The receive weights *ω*_*q*_ include the normalized angular quadrature and any prescribed apodization; uniform unapodized sampling uses *ω*_*q*_ = 1*/N*_rx_. The retained mode set is N. Thus 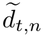 is the spectrum of one angular pattern.

For propagation in these coordinates, we construct a periodic reference kernel whose angular dependence is a convolution. Its Fourier modes propagate independently, as proved in Appendix A-I. This diagonal relation is a property of the reference, while (8) supplies modal coordinates for the measured finite-arc data. We next specify the radial phase of that reference.

For a mode 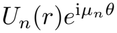, separation of the homogeneous polar Helmholtz equation gives

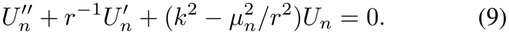

On the outgoing propagating branch, write the leading oscillatory field as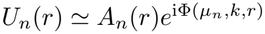, with slowly varying amplitude *A*_*n*_. The radial phase derivative is 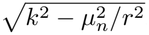 Integrating it gives the radial phase

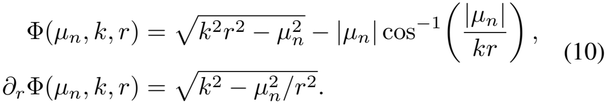

Appendix A-II derives this phase from the radial equation and its outgoing branch.

Choose a reference transmit path 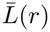 that depends only on radius. Its transmit phase is 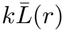, while outward receive propagation contributes Φ(*µ*_*n*_, *k, r*) − Φ(*µ*_*n*_, *k, r*_0_). Their sum is

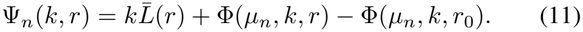

The coefficient 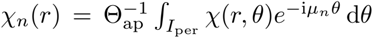 projects the angular object distribution at radius *r* onto the *n*th Fourier pattern. Division by Θ_ap_ gives the normalized coefficient used in Appendix A-I. The corresponding phase-form modal reference is

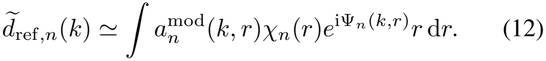

Here 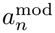 collects forward source and propagation amplitudes, and subscript ref identifies the selected radial-transmit reference. A centered source has 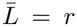. The propagating-phase derivation requires *kr >* |*µ*_*n*_| and *kr*_0_ ≥ (1 + *σ*) |*µ*_*n*_|, with margin *σ >* 0 at the receive radius. The semicircular implementation extends the reference anchor by clipping outside this latter branch; its exact convention and local support are specified in Supplementary Section S4.1.

### B. Local Stolt Mapping and Reference Reconstruction

The radial phase in (12) is nonlinear in *r*, whereas radial Fourier synthesis requires a phase linear in the output coordinate. We therefore partition the image radius into overlapping intervals and linearize the phase within each one. On *I*_*j*_ = [*r*_*j*_ − *h*_*j*_, *r*_*j*_ + *h*_*j*_], with center *r*_*j*_ and half-width *h*_*j*_,

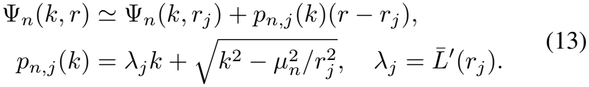

For angular mode *n* in interval *j, p*_*n,j*_(*k*) = *∂*_*r*_Ψ_*n*_(*k, r*_*j*_) is the radial phase slope at the interval center, with units of inverse length. Stolt remapping uses this slope as the radial Fourier coordinate. When *λ*_*j*_ = 1, as for 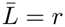 the inverse and its Jacobian are

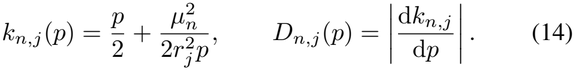

The retained *k* band maps monotonically to *P*_*n,j*_ on the propagating branch; for (14), this branch has *p >* | *µ*_*n*_| */r*_*j*_. Supplementary Section S2 gives the inverse for any *λ*_*j*_ *>* 0.

The reference phase now supplies a migration rule. We reverse it, select the propagating support with *W*_*n,j*_, and apply a fixed image filter 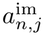. Changing variables from *k* to *p* gives

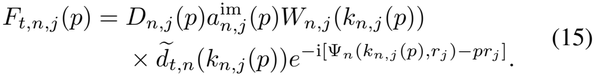

The Jacobian *D*_*n,j*_ compensates the change in spectral sampling density through d*k* = *D*_*n,j*_d*p*, preserving the integral under remapping. The exponential removes the local phase intercept, leaving *e*^*−*i*pr*^ for radial synthesis. The image filter 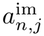 controls spectral weighting; it is chosen separately from the forward amplitude 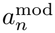 in (12). Its implemented regularized form is given in Supplementary Section S4.1. Together these choices define the migration operator used below.

For a source at the curvature center, denoted by subscript 0, inverse radial Fourier synthesis and angular summation give a field on each interval. Their weighted overlap forms the reference image:

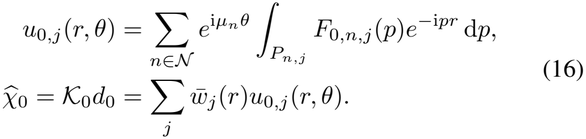

The integral reconstructs each radial modal field, the angular sum forms the local image *u*_0,*j*_, and the interval sum defines _0_. Subscript 0 denotes the centered transmit event. Nonnegative overlap windows have positive total weight over the image and are normalized as 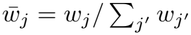. The implemented square-root-Hann windows blend neighboring local reconstructions. Offset transmissions use this same synthesis structure with an additional path correction.

### C. Offset-Phase Factorization and Image Formation

The reference reconstruction uses a transmit path that depends only on radius. For an offset source, points on the same radius have different transmit distances. Their phase correction must be included before coherent compounding. With the receivers fixed as in Fig. 1, the additional path for transmission *t* is

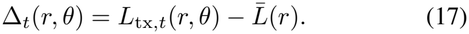

This path introduces a position-dependent correction inside the radial integral. Reversing its phase in the reference synthesis gives

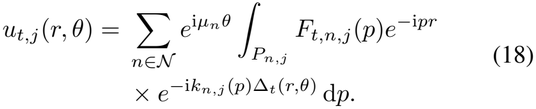

The offset factor depends jointly on mode *n*, radial wavenumber *p*, and output position, preventing its direct inclusion in a position-independent spectrum. We first extract the part that changes the radial sampling coordinate, then separate the residual into reusable spectral factors and spatial weights. The unit-reference-slope inverse in (14) illustrates the distinction:

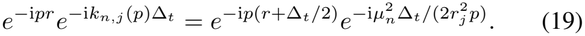

The first factor changes the query radius by Δ_*t*_*/*2; the second remains dependent on mode and frequency. A *radial pullback* evaluates a synthesized complex field at a geometry-dependent radius for each output pixel. The following construction chooses this query over the retained support and represents the remaining phase with screens.

For the following local construction, fix a transmit event *t* and interval *j* and suppress these two indices. In particular, *F*_*n*_ denotes the weighted spectrum *F*_*t,n,j*_ in (15), whose centered-transmit instance *F*_0,*n,j*_ supplies (16). Write *k*_*n*_(*p*) = *ηp* + [*k*_*n*_(*p*) − *ηp*]. A single slope *η* is chosen across retained modes to minimize the range of the spectral residual *k*_*n*_(*p*) − *ηp*. We also subtract a constant path center *ℓ*^*c*^ from Δ: centering the path reduces the variation left for the phase screens. Without a query-domain constraint, this center is the midpoint of the path extrema in the interval and imaging sector. The term *ηp*(Δ− *ℓ*^*c*^) can then be combined with *pr*, defining the query radius

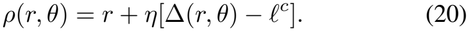

The field is read at (*ρ*(*r, θ*), *θ*) to form the output at (*r, θ*). For a fixed event, interval, and output pixel, all modes and expansion terms share this query radius. It varies across the image and, in general, with *t, j*. The center *ℓ*^*c*^ is a path reference for this decomposition; its admissible values also keep *ρ* inside the prepared synthesis domain. Supplementary Section S3 gives the construction of *η* and the constrained center.

Within the coupled target (18), this change gives the exact identity

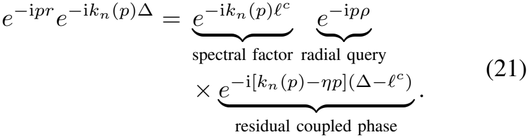

The identity retains both the spectral path-center factor and the residual coupled phase. To evaluate the latter through synthesized fields, we combine it with the spectral factor in a finite Chebyshev–Bessel expansion [35]:

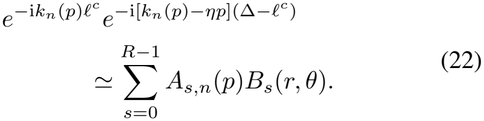

Here *R* is the number of retained terms. The spectral factor *A*_*s,n*_ multiplies the data before synthesis, while the spatial screen *B*_*s*_ multiplies the synthesized field. Each *B*_*s*_ is a complex amplitude-and-phase weight independent of *n, p*. Appendix B constructs both factors and derives the image formula.

Substituting (22) into the coupled target (18) therefore reduces its evaluation to fields defined by

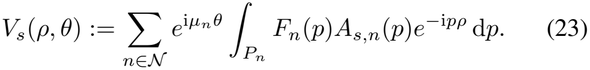

Because *A*_*s,n*_ is independent of output position, each field is synthesized from the channel spectra once per transmit event and interval over its query domain. Restoring *t, j*, sampling these fields and blending the intervals defines the finite-term image:

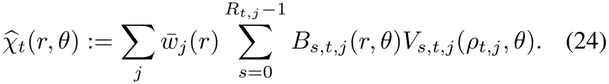

The map 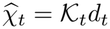 approximates the interval-weighted target through the truncation in (22). Equations (23) and (24) define its computed fields and output; Supplementary Section S3.1 bounds the truncation. The convex example uses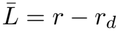, the exact path on the source ray, so its unit radial slope gives (14).

Dropping the residual coupled factor leaves an affine phase approximation, whose error envelope is evaluated in Section IV-B. The following scalar-pullback construction fits a geometrical displacement of a reference image.

### D. Scalar-Pullback Approximation

The phase-screen construction retains the residual coupling displayed in (19) through several synthesized fields. A simpler option is to represent the transmit correction by a displacement of one reference image. A common affine inverse Stolt map permits an exact shift and spatial carrier; the nonaffine maps here require an approximation. Scalar pullback defines this displacement through a geometrical path fit. This choice approximates the offset correction at the query-construction stage, while *R* in (22) controls the subsequent spectral factorization. We specify the semicircular realization first and then its exact-path counterpart used in the convex comparison.

For this realization, take 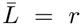 and retain the second-order expansion

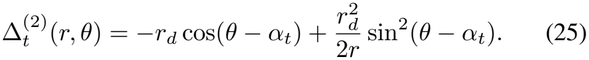

The expansion is in *r*_*d*_*/r*, with path remainder 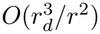. Its angle-independent term 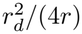 follows from the identity sin^2^ *ϑ* = (1 − cos 2*ϑ*)*/*2.

Let 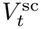 be the reference image before scalar-pullback correction, with superscript sc identifying this implementation. In the semicircular example, it combines radial modal fields using mode-dependent normalized overlap and angular summation. The angle-independent path term changes the transmit slope to 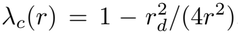. The implemented Stolt map uses this slope, but its phase at *r*_*j*_ remains the centered-source value. This combination introduces a linear contribution to the local phase residual, in addition to the usual Taylor remainder. Supplementary Section S2 defines the reference field, and Section IV-B evaluates this residual.

To convert a transmit-path increment into a radial displacement, define the total geometrical path slope 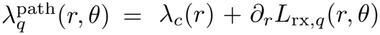. At each radius, the scalar coefficient Minimizes 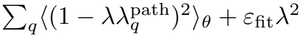, giving

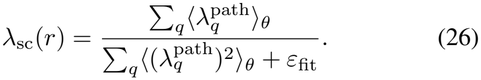

Here ⟨·⟩ _*θ*_ is the uniform mean over the image-angle grid, and *ε*_fit_ *>* 0 stabilizes the denominator. The fit uses geometry alone and is shared by transmissions at the same source radius.

For the semicircular path expansion, its resulting query is

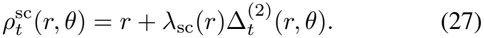

The single-transmit image is obtained by one radial pullback of the reference image:

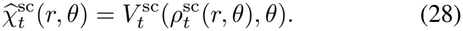

Equation (28) replaces the sum of screen-weighted fields in by one evaluation of 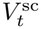. This reduces synthesis work at the cost of the scalar phase approximation assessed in Section IV-B. Supplementary Section S1 provides starting configurations and a worked accuracy example.

The convex scalar control uses the same reference 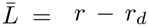 and exact transmit distance as the convex phase-screen implementation. Its geometrical fit uses a reference transmit slope of one, normalized receive quadrature, and trapezoidal averaging over the image-angle grid. In (27), 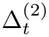 is replaced by the exact residual *L*_tx,*t*_ − (*r* − *r*_*d*_); (28) then samples the corresponding reference image. This common-path comparison introduces no transmit-path expansion error. The semicircular comparison retains its established reference synthesis and adds an exact-path phase-screen evaluation. Supplementary Section S6 specifies both numerical realizations and their controls.

### E. Coherent Compounding

Both implementations now provide a complex image for each transmission. Denoting either output by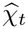, we combine them with prescribed nonnegative weights *v*_*t*_(*r, θ*), normalized to Σ_*t*_ *v*_*t*_ = 1:

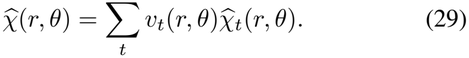

Envelope detection and logarithmic compression follow this coherent sum, preserving relative transmit phases during compounding. At zero offset, both constructions return their reference synthesis. Geometry-dependent factors and transform plans are reusable, while the fields and their sum are evaluated for each RF acquisition; Supplementary Section S6.2 gives the computational costs [36].

The next section first evaluates the local and offset-phase approximations, then compares DAS and both CAMM implementations on the same point-target RF within each geometry.

Part II illustrates the numerical workflow with RF spectra and intermediate images.

## IV. Error Control and Numerical Validation

The validation connects the approximation controls to the resulting point-target responses. After specifying the simulations, we examine the phase-screen implementation and then the scalar-pullback approximation, following Section III. Within each geometry, the same point-target RF is reconstructed by DAS, CAMM phase screen, and CAMM scalar pullback. A direct-quadrature check of the separated numerical evaluation is reported in Supplementary Section S5.

### A. Validation Setup

Table II gives two outward-looking circular-arc configurations with contrasting array and image scales. The compact semicircular array provides a wide angular aperture with small source-to-target radius ratios. The conventional convex array uses substantially larger relative offsets. Comparing both CAMM implementations in each configuration tests whether a scalar correction preserves the response obtained with phase screens.

**TABLE II.** Probe and Transmit Parameters.

| Parameter | Semicircular | Convex array |
|---|---|---|
| Background sound speed *c*<sub>0</sub> | 1540 m/s | 1540 m/s |
| Radius *r*<sub>0</sub>/*r*<sub>*d*</sub> (mm) | 0.75 / 0.375 | 55 / 27.5 |
| Receive elements / TX events | 64 / 24 | 192 / 24 |
| Receive aperture | 180° | 70° |
| Angular spacing | 2.8125° | 0.364583° |
| First / last receive angle | −90° /<br>87.1875° | −34.817708° /<br>34.817708° |
| Uniform source-angle range | ±90° | ±35° |
| Center frequency (MHz) | 20 | 3.5 |
| Nominal pulse bandwidth | 40% | 65% |
| RF sample rate (MHz) | 200 | 100 |
| Samples per channel | 2225 | 14034 |

Both use offset virtual sources and coherent compounding. Pulse bandwidth is fractional; the retained processing bands are 8–24 MHz and approximately 2.37–4.64 MHz, respectively. The additional isolated convex pair uses 16000 samples per channel with the same probe and transmit parameters.

The point-target comparisons in Section IV-C use rectangular receive apodization (equal element weights) and no additional modal taper. Within each geometry, DAS and CAMM share the RF band and normalized trapezoidal transmit weights: *v*_1_ = *v*_24_ = 1*/*46 and *v*_*t*_ = 1*/*23 otherwise. The convex RF band has a cosine taper over 10% of its samples in total, split equally between the two edges and applied identically to both methods. CAMM retains the Stolt density *D*_*n,j*_, propagating-support conditioning *W*_*n,j*_, normalized square-root-Hann overlap windows, and the regularized migration amplitude 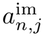 specified in Supplementary Section S4.1. These factors are fixed before RF processing. Controlled reconstructions vary the receive/modal taper and post-Stolt amplitude separately, with the phase operations unchanged.

#### Semicircular configuration

The phantom contains 20 targets at all combinations of *r* ∈{2.5, 3.5, 4.5, 5.5} mm and *θ*∈{− 60, − 30, 0, 30, 60} ^*°*^. All three reconstructions use a 260× 400 grid over *r* = 1.5–7 mm and *θ* ∈ [−90^*°*^, 90^*°*^). DAS uses analytic RF filtered to the same 713 bins over 8–24 MHz; all methods use the 24 transmissions and original time origin. The phase-screen calculation retains the semicircular spectral reference, including its clipped receive anchor, and evaluates the exact offset path with separated factors. Scalar pullback uses (25)–(28). Peaks are searched on the reconstruction grid within ± 0.4 mm and ± 10^*°*^ of each target. Tests of the reconstruction coefficients use a 66 × 101 grid with the same band and local propagating support among 61 candidate modes, without inserting channel-data values.

#### Convex configuration

Four targets lie at (*x, z*) = (0, 10), (− 18, 30), (18, 50), and (0, 70) mm, where *z* is depth from the array apex. The 256 × 257 reconstruction grid spans *r* −*r*_0_ = 5–75 mm and *θ* = ± 27.5^*°*^. The phase-screen calculation uses 19 radial intervals, 638 frequency bins over 2.3657–4.6352 MHz, and 191 candidate modes. The scalar control uses the same exact transmit path and source-ray reference, with a geometrically fitted query. All three methods share RF, frequency conditioning, and all 24 transmissions. The radial-phase tolerance is 0.3 rad. An additional pair of isolated points compares *θ* = 0^*°*^ and − 20^*°*^ at the same radial depth *r* − *r*_0_ = 10 mm, using the same array, excitation, and output grid. The matched-test protocol and point-response metrics for this pair are given in Supplementary Section S6.1. The CAMM comparisons retain each geometry’s reference convention and common acquisition settings. Their radial synthesis grids, interval blending, and interpolation follow the respective implementations. Thus the point responses compare complete implementations; the component tests above and in the supplement examine individual approximation steps. Full query coverage and phase-screen convergence are checked before measuring the images (Supplementary Sections S5 and S6).

### B. Approximation Errors

The phase-screen implementation retains the exact transmit path. Its approximation controls therefore concern the local radial reference phase and truncation of the separated off-set factor. The scalar-pullback tests additionally examine the transmit-path expansion and replacement of the offset phase by one radial query.

#### Radial linearization

Here the local phase is the consistent Taylor approximation in (13). Its largest departure from (11) is

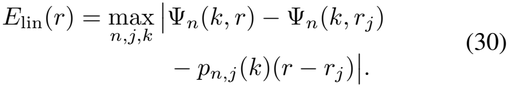

The subscript lin denotes radial linearization. The maximum is over retained native mode–frequency pairs and active intervals with *w*_*j*_(*r*) *>* 0. This quantity, in radians, tests whether each interval is short enough for the local phase approximation. For the convex configuration, its maximum is 0.2722 rad, below the prescribed 0.3-rad tolerance. The Taylor bound and radial-error summary are given in Supplementary Sections S3.1 and S5.

#### Offset phase left by an affine correction

The factorization (21) and Appendix B leave the coupled factor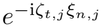, With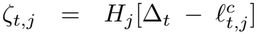, spectral-residual half-width *H*_*j*_, and |*ξ*_*n,j*_| ≤1. Its phase envelope is *E*_aff_ (*r, θ*) = max_*t,j*_ |*ζ*_*t,j*_(*r, θ*) |, restricted to active intervals. Figure 4(a) locates its maximum of 9.286 rad in the shallow off-axis region. An affine-only evaluation replaces this coupled factor by unity; panel (c) bounds the resulting complex-factor discrepancy. Its upper limit of two corresponds to unit-magnitude factors with opposite phases. These panels identify where a coordinate shift and carriers leave substantial phase variation for the screens to represent.

**Fig. 4.**
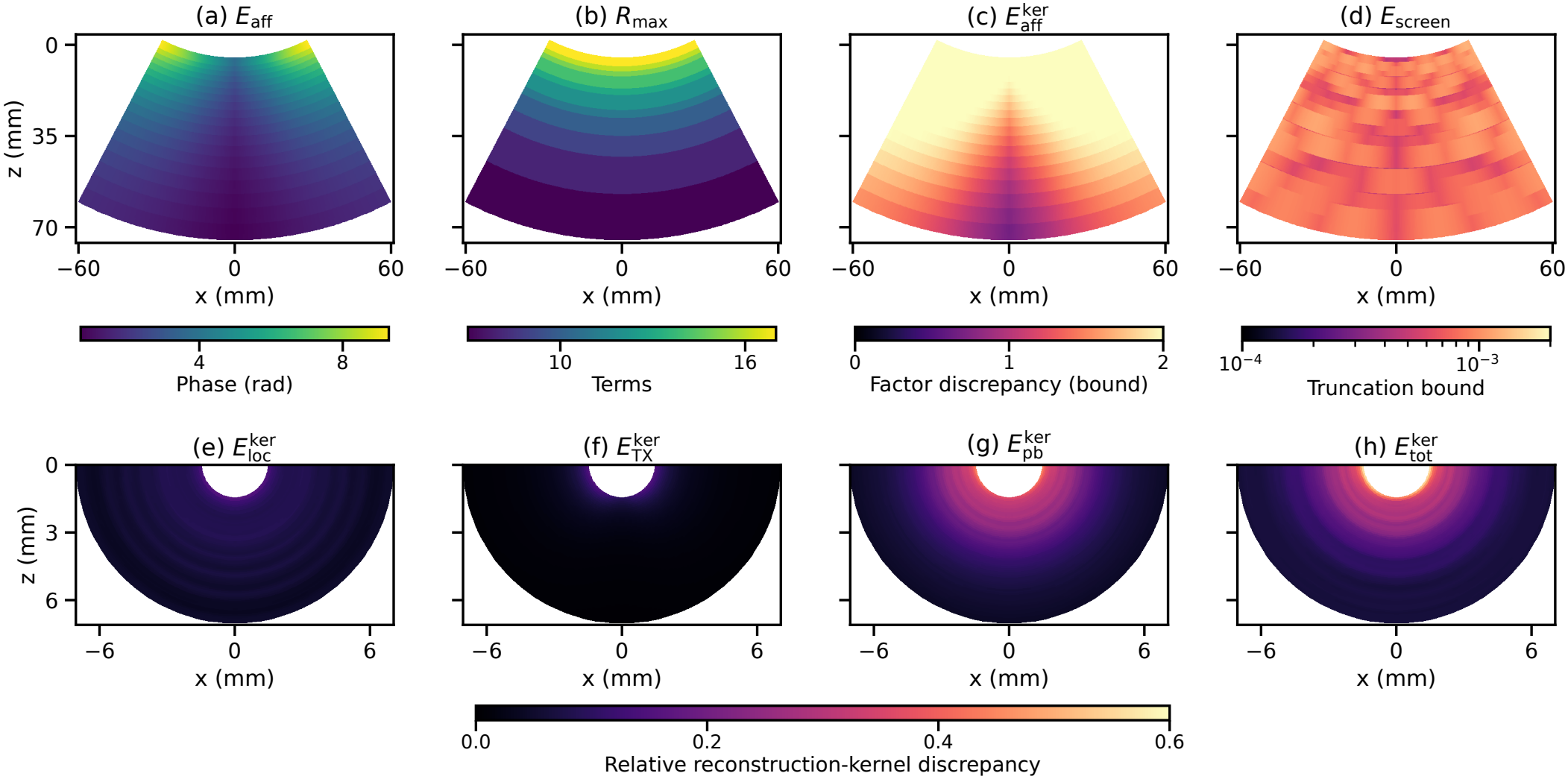
Spatial approximation controls. Convex phase-screen controls: (a) residual-phase envelope, (b) retained terms, (c) affine-only complex-factor envelope, (d) truncation bound. Semicircular reconstruction-coefficient discrepancies: (e) local synthesis, (f) transmit-path expansion, (g) scalar pullback, (h) combined phase. Panels (e)–(h) share a scale; (c,d) have separate, labeled scales.

#### Phase-screen truncation

Keeping *R*_*t,j*_ terms in (22) leaves the Bessel-series tail from (B.2). Over the same active intervals, the complex-factor error is bounded by

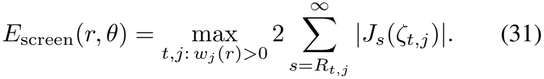

Here *J*_*s*_ is the Bessel function in (B.2); screen identifies truncation of the phase-screen series. This dimensionless bound limits the complex-factor difference, complementing the phase-angle bound in (30). Retaining 4–17 terms [Fig. 4(b)] gives a maximum bound of 1.243 × 10^*−*3^ [panel (d)]. The largest directly sampled residual-factor error is 9.914× 10^*−*4^, within that bound. The higher term counts in the shallow off-axis region therefore control the stronger coupling seen in panel (a). Panels (c,d) use separate color scales to show the spatial variation within their different error ranges. Supplementary Sections S3.1 and S4.3 give the proof and map definitions.

#### Scalar-pullback approximation

This implementation replaces the offset correction by a displacement of one reference image. To distinguish its approximation steps, we compare reconstruction coefficients without inserting channel-data spectra. These tests therefore describe the reconstruction rule independently of the particular simulated target arrangement. At pixel **x**, let **h**_test_ collect the complex coefficients multiplying the retained mode–frequency–transmit samples. They form the discrete synthesis kernel for that pixel. For each test below, the specified signed phase residual is inserted before combining overlapping intervals; **h**_ref_ uses the same support and weights with that residual set to zero. For ∥ **h**_ref_ ∥ _2_ *>* 0, define the relative reconstruction-coefficient discrepancy

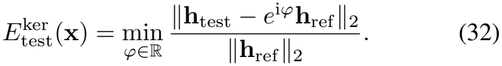

The superscript ker identifies the synthesis kernel. One fitted phase *Φ* per pixel is shared across all modes, frequencies, and transmissions. It removes a common rotation of the compounded output, which leaves the envelope unchanged; zero means coefficient agreement up to that rotation. Supplementary Sections S2.2 and S4.2 give the signed residuals and coefficients.

Figure 4(e)–(h) applies (32) to four tests in the semicircular configuration. Their labels identify the operation being assessed.

#### Local synthesis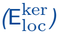

This test compares the phase used to synthesize 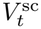 with the unlinearized centered-source phase at the query radius. It includes the corrected slope and centered anchor described in Section III-D. Panel (e) varies with the active intervals and has maximum discrepancy 0.160. In the refinement test, the anchor–slope combination contributes a term linear in query-to-center distance, whereas the consistent Taylor control has a quadratic remainder. Supplementary Section S5 summarizes their different refinement behavior.

#### Transmit-path expansion (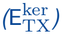)

Replacing the exact path increment (17) by (25) changes the transmit phase by the path remainder multiplied by *k*. Panel (f), with maximum 0.151, concentrates this discrepancy near the inner radius, where *r*_*d*_*/r* is largest. The source-radius refinement test approaches the cubic remainder predicted by the second-order expansion.

#### Scalar pullback (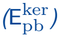)

The abbreviation pb denotes pull-back. Holding the second-order path fixed, this test compares its use as the offset factor in (18) with the radial displacement in (27)–(28). Panel (g) isolates the replacement of a mode– frequency-dependent correction by one coordinate shift. It gives the largest individual shallow discrepancy, with maximum 0.441, and its residual persists when the radial partition is refined.

#### Combined phase (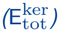)

We add the three signed phase residuals before evaluating (32). Panel (h) has maximum 0.557 and median 0.118. Evaluating the combined phase retains reinforcement and cancellation among the three residuals.

At the nominal target radii 2.5, 3.5, 4.5, and 5.5 mm, the maximum combined coefficient discrepancy over the five target angles is 0.294, 0.182, 0.111, and 0.079, respectively.

Its decrease with radius locates the larger scalar-phase changes at shallow positions. We next examine the reconstructed point responses at these positions.

### C. Point-Target Imaging

For the tested semicircular array, scalar pullback preserves the principal point response obtained with phase screens. In Fig. 5, both CAMM implementations retain the twenty targets and have less conspicuous arc-shaped responses than DAS, although circular residual structure remains. Their four angular cuts resolve the same five peaks per ring. All twenty CAMM radial peak indices coincide; fourteen angular peak indices coincide and the remaining six differ by one 0.45^*°*^ angular pixel. The maximum angular deviation is 0.30^*°*^ for either implementation, compared with 1.65^*°*^ for DAS (Table III).

**TABLE III.** Point-target localization and envelope width for the three reconstruction methods.

| Geometry | Quantity | DAS | CAMM phase screen | CAMM scalar pullback |
|---|---|---|---|---|
| Semicircular (20 targets) | Max. angular deviation (°) | 1.65 | 0.30 | 0.30 |
|  | Radial displacement (mm) | 0.162–0.168 | 0.162–0.168 | 0.162–0.168 |
|  | Position error (mm) | 0.170–0.200 | 0.162–0.168 | 0.162–0.168 |
|  | Radial FWHM (mm) | 0.113–0.118 | 0.114–0.117 | 0.117–0.122 |
|  | Tangential FWHM (mm) | 0.138–0.459 | 0.155–0.483 | 0.158–0.494 |
| Convex (4 targets) | Position error (mm) | 0.059–0.178 | 0.059–0.178 | 0.059–0.178 |
|  | Radial FWHM (mm) | 0.587–0.677 | 0.637–0.682 | 0.625–0.673 |
|  | Tangential FWHM (mm) | 0.276–0.687 | 0.430–0.906 | 0.532–0.914 |

Ranges are across the targets in Figs. 5 and 6. Position error is the Euclidean distance from the nominal target to its detected peak; radial displacement is signed. FWHM uses the nearest interpolated half-amplitude crossings around that peak. Tangential width follows the detected radial arc.

**Fig. 5.**
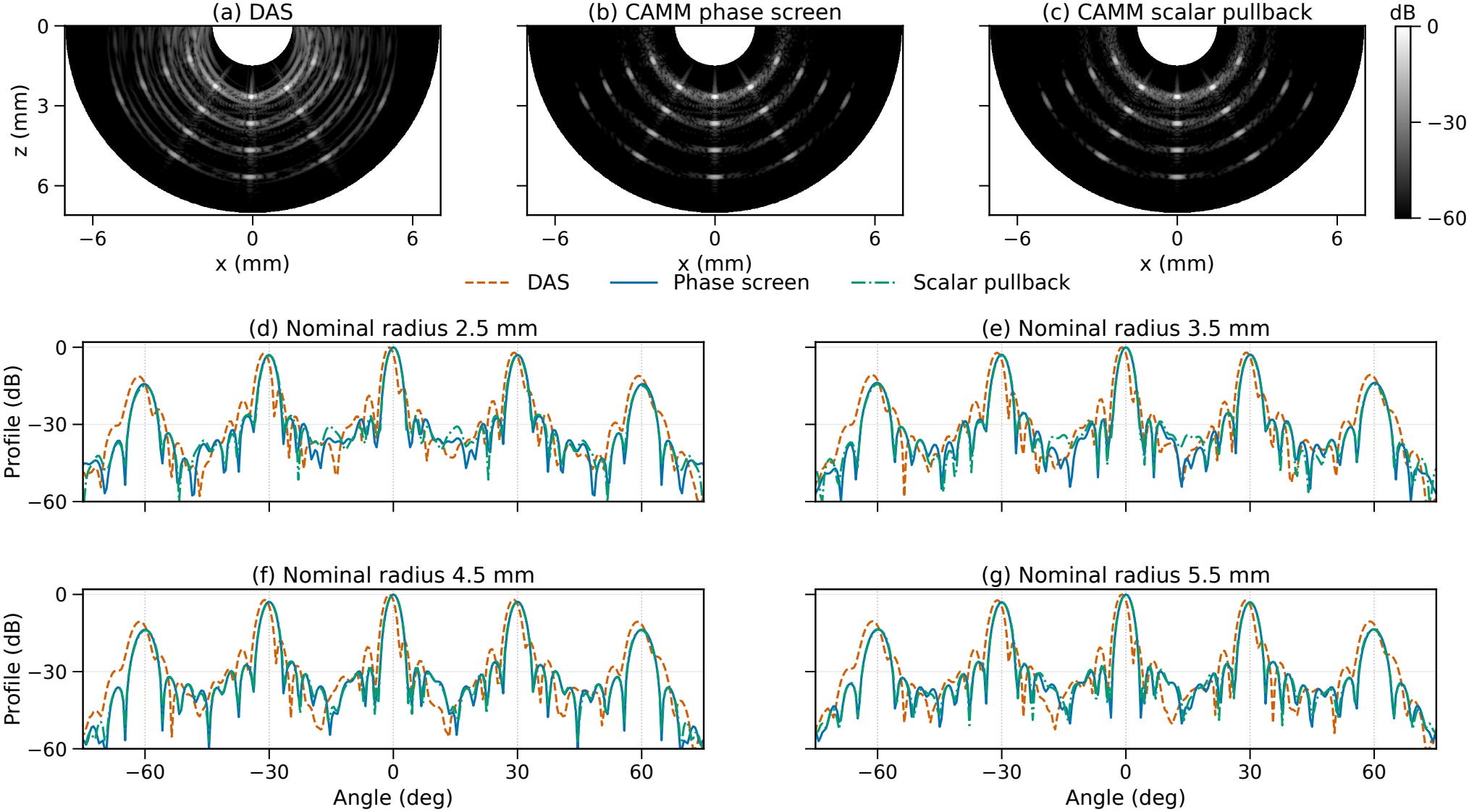
Same-RF semicircular comparison. (a)–(c) DAS, CAMM phase screen, and CAMM scalar pullback. (d)–(g) Angular profiles at each method’s median detected radius for the indicated target ring; dotted lines mark nominal angles. Images and profiles use whole-field and per-cut peak normalization, respectively, over 60 dB. Receive weights are rectangular and the additional modal taper is unity.

Scalar pullback gives median radial and tangential FWHM values of 0.119 and 0.316 mm, compared with 0.116 and 0.309 mm for phase screens. The small width changes support using the simpler scalar realization when these point-response differences meet the imaging requirement. Phase-screen evaluation also changes the surrounding response and amplitude, which remain part of that assessment. All three methods retain the 0.162–0.168-mm radial displacement; widths are measured around their detected peaks in the original image coordinates. The matched modal point-response measurements in Supplementary Section S1.1 help interpret these width changes: the interpolation-based scalar evaluation adds radial broadening and attenuation relative to its phase proxy. The channel-RF comparison includes this numerical interpolation as part of the scalar implementation.

The convex comparison makes the need to inspect more than the peak position particularly clear. All three methods locate the four targets in the same grid cells, but the scalar image has more pronounced cross-shaped and radial responses around the peaks in Fig. 6. These structures remain visible in the full sector and in the target-centered views, which retain the whole-image normalization. Phase-screen evaluation reduces these structures while retaining the exact transmit path used by the scalar control.

**Fig. 6.**
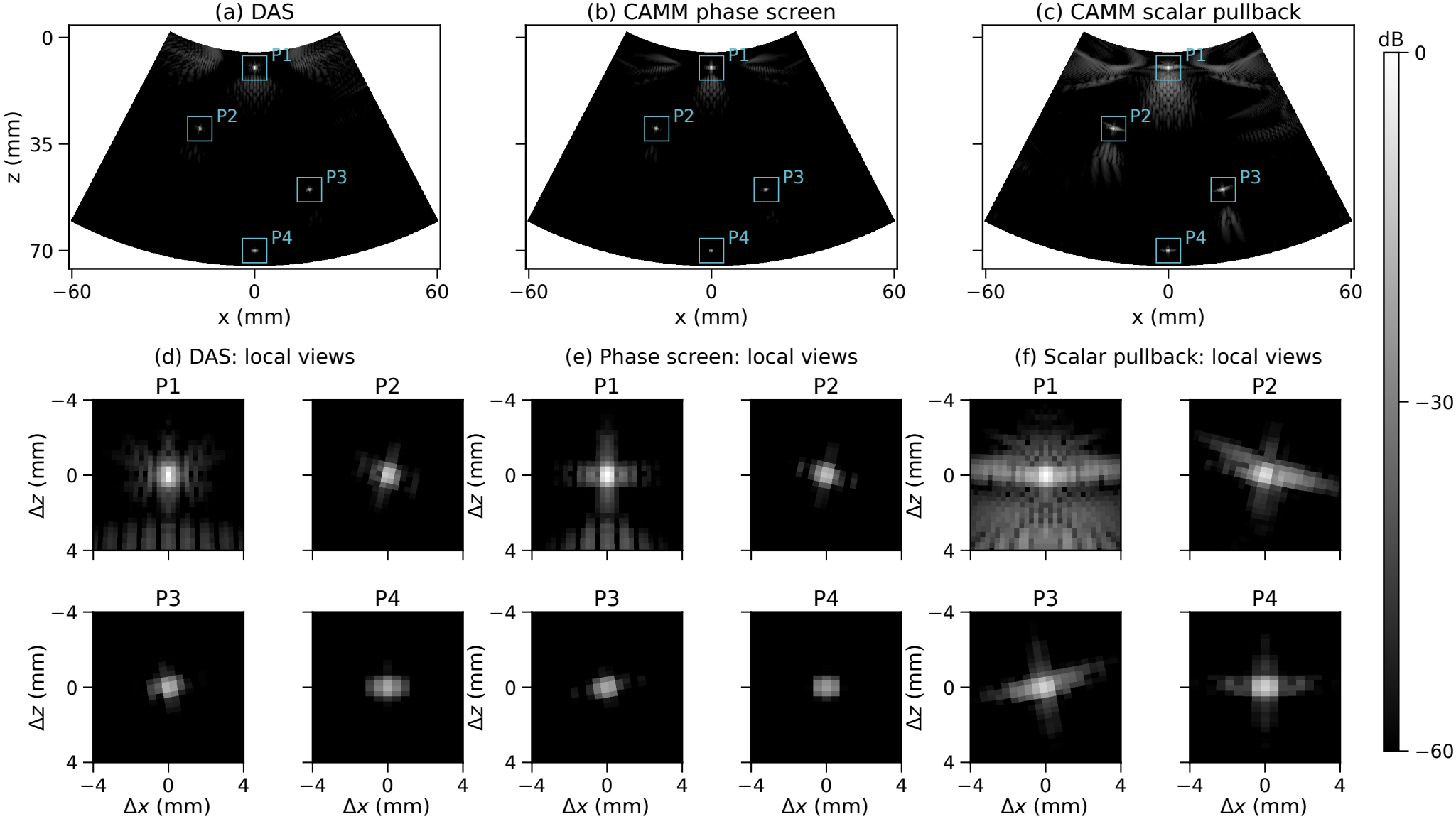
Same-RF convex comparison. Columns show DAS, CAMM phase screen, and CAMM scalar pullback. (a)–(c) Full fields; (d)–(f) the corresponding four target-centered views, ordered P1–P4 by depth. Every 8 *×* 8 mm local view retains its method’s whole-image normalization over 60 dB. Both CAMM implementations use the exact transmit path.

At the shallow axial target P1, tangential FWHM is 0.532 mm with scalar pullback and 0.430 mm with phase screens. The corresponding values are 0.714 and 0.679 mm at P2, 0.780 and 0.823 mm at P3, and 0.914 and 0.906 mm at P4. At P3, the scalar central lobe is slightly narrower and its surrounding response more extended. DAS has the narrowest tangential lobes at all four targets (Table III). The main-lobe width and the surrounding structure therefore answer different questions. Together with the residual-phase maps in Fig. 4(a)–(d), the full images support retaining phase screens when surrounding-response control is required in this convex configuration.

The isolated shallow pair quantifies this surrounding response without contributions from other targets. We measure the fraction of full-field squared image magnitude outside a fixed neighborhood extending ±1 mm radially and ±3 mm tangentially around the known target, using polar pixel-area weights. At 0^*°*^, this fraction is 4.14% for DAS, 1.36% for phase screens, and 13.47% for scalar pullback; at − 20^*°*^, it is 3.45%, 0.61%, and 11.82%, respectively. All three methods retain the same peak cell within each case. These paired responses support the phase-screen choice for controlling extended energy in this shallow convex setting, separately from the ranking of central-lobe widths.

Figure 7 complements these energy fractions with the isolated off-axis images and their output angular spectra, computed by angular Fourier transformation of the completed complex images.

**Fig. 7.**
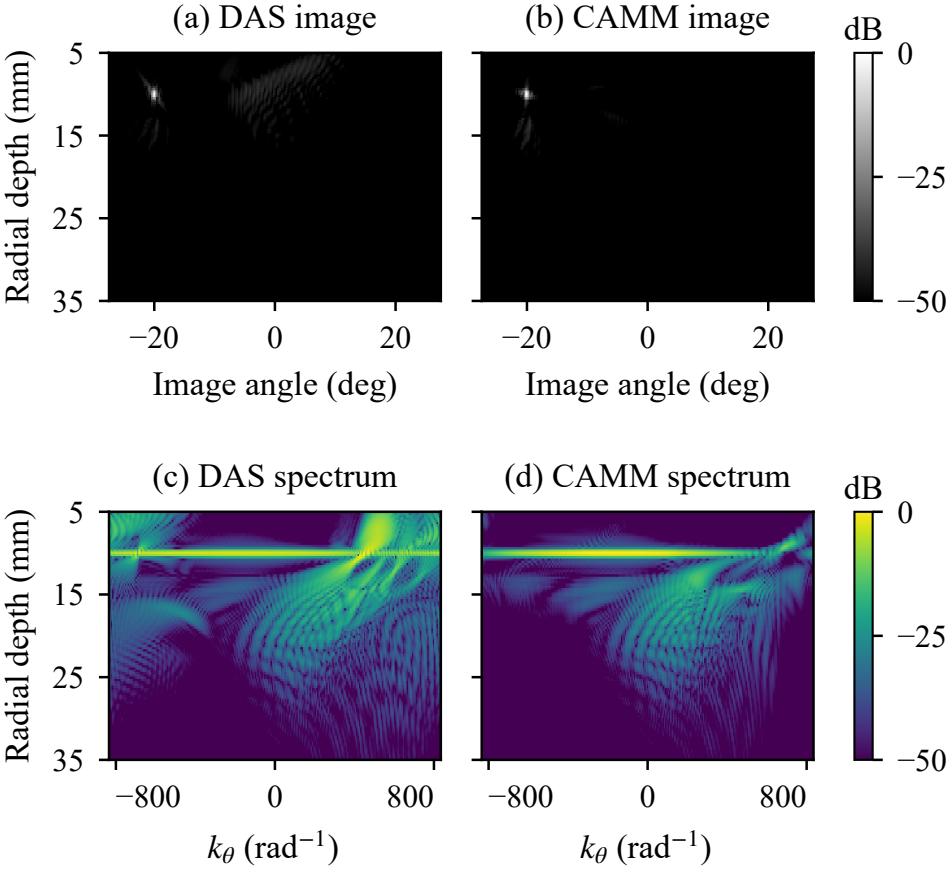
Isolated point at radial depth 10 mm and angle *−* 20^*°*^. (a), (b) DAS and CAMM phase-screen images; (c), (d) corresponding output angular amplitude spectra versus angular spatial frequency *k*_*θ*_. The same full-angle Hann window precedes each angular FFT. Each image and spectrum uses its own full-grid peak and a *−* 50 to 0 dB display range, showing relative response structure.

The one-factor reconstructions in Supplementary Section S5.1 test additional contributions to response shape in the semicircular scalar and convex phase-screen baselines. Compared with the Lanczos-tapered control, the untapered semicircular baseline has narrower tangential lobes at unchanged peak cells. Convex receive-taper and unit post-Stolt multiplier controls leave the principal widths nearly unchanged, whereas the semicircular multiplier control changes weak outer responses more strongly. These controls and the matched modal interpolation test complement the complete three-method comparisons.

## V . Discussion

The error-control results establish the numerical feasibility of the CAMM construction. Radial subdivision and phase-screen tail bounds control the numerical approximations, while agreement with direct integration verifies the separated evaluation of the selected modal imaging integral. The scalar tests assess the further approximation of the offset correction. Together, these checks connect the propagation model to an accuracy-controlled synthesis of coherent images from channel spectra.

With this numerical basis established, the phase-screen images show weaker arc-shaped responses in the semicircular array and weaker slanted structures around the shallow off-axis convex target. Figure 7(a), (b) illustrates the latter change while retaining the target response. In (c), (d), the main spectral band remains near the target depth, while structures at other depths are relatively weaker under CAMM. Here *k*_*θ*_ is the angular spatial frequency of the completed complex image. The separately peak-normalized spectra illustrate relative response structure, while the preceding energy fractions quantify its spatial distribution. For this convex target, the weaker surrounding response accompanies a wider central tangential lobe, revealing a tradeoff between response concentration and central-peak resolution.

This redistribution can be understood through the coherent reinforcement of contributions away from the target. Physical angular frequencies separated by 2*π/*Δ*β*, with receive-angle spacing Δ*β*, have the same uniformly sampled pattern up to a constant phase. Such sampling matches can let the DAS delay sum reinforce contributions at a location away from the true point. CAMM synthesizes each retained order *µ*_*n*_ with its radial phase in (10). This changes the relative phase across frequency and mode and can weaken the resulting off-target reinforcement. For a sampling replica and the retained order on the same propagating branch, the radial phase slopes differ by

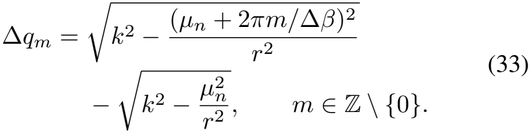

Here *m* identifies a sampling replica. The expression describes interior propagating contributions, with both orders remaining separated from their turning points along the radial path. Equation (33) gives the rate at which their radial-phase mismatch accumulates. When the resulting relative phase variation across frequency and mode weakens an off-target coherent sum, its amplitude decreases. This mechanism connects the modal propagation assignment to the weaker surrounding responses observed in the tested configuration.

For a finite arc, target angle changes the propagation directions sampled by the aperture and their contributions to such off-target matches. Phase alignment and amplitude weighting therefore jointly shape the position-dependent image response illustrated in Fig. 7. Within CAMM, phase screens preserve the spatially varying offset-transmit coupling during this synthesis. The structure of this coupling also determines when the offset correction can be simplified to one radial query while preserving the required response. In (13), | *µ*_*n*_| */*(*kr*_*j*_) compares tangential spatial frequency with acoustic wavenumber. Similar normalized radial slopes *p*_*n,j*_*/k* across contributing modes and frequencies favor this scalar approximation. Their variation leaves the coupled phase in (19), whose magnitude also depends on the transmit-path increment. Modal content, bandwidth, image radius, and source displacement therefore determine the choice together.

The compact high-frequency semicircular array illustrates a useful scalar approximation: it closely preserves the principal widths and peak positions of the phase-screen image. In the convex examples, scalar pullback retains the detected positions and produces more pronounced cross-shaped and radial responses. The implementation choice follows the required central and surrounding response. Supplementary Table S1 provides engineering starting points. For an unlisted acquisition, the frequency band, relative source displacement, and angular spectrum guide an initial choice, followed by comparisons at representative shallow, off-axis, and deeper positions. The supplementary kernel scans and point-response tests provide examples for checking a chosen accuracy requirement.

This choice has a direct computational consequence. Each retained screen term adds field synthesis and spatial accumulation, whereas scalar pullback reads one reference image. A satisfactory scalar approximation can therefore reduce this work. Geometry-dependent factors and transform plans can be reused at fixed acquisition and image grids; data-dependent fields still require computation. Part II assesses how implementation and hardware affect timing and storage alongside image quality for the evaluated reconstruction pipelines.

The radial propagation relation also connects CAMM to linear-array spectral imaging. As the curvature radius tends to infinity with physical aperture and local coordinates fixed, angular modes become transverse Fourier components and the radial phase approaches Cartesian propagation. With a corresponding transmit reference, the local planar limit in Supplementary Section S3.2 recovers Stolt-type dispersion [23], [25]. A displaced virtual source can still leave a spatially varying transmit path. Its treatment remains an accuracy– computation choice: phase screens retain residual coupling through weighted fields, while scalar pullback approximates it through a radial query. Both approximate the same modal migration target.

## VI. Conclusion

CAMM establishes a direct spectral route from circulararc channel data to coherent images, extending Stolt-type migration to radius-dependent modal propagation and spatially varying offset-transmit phase. Its central contribution is a common modal construction that integrates radial spectral synthesis with offset-phase compensation. Phase-screen factorization retains the coupled phase through spectral factors and spatially weighted fields, while scalar pullback provides a compact approximation within the same construction. These realizations unify the treatment of semicircular and convex-array configurations and provide an explicit accuracy–computation choice.

Error controls and direct-integral verification connect the propagation model to its numerical realization, while matched-data point-target studies demonstrate its influence on central and surrounding responses. The local planar limit further connects the framework to established linear-array spectral migration. Together, these contributions establish a systematic basis for geometry-adapted spectral image formation on circular arcs, linking propagation physics, approximation accuracy, and reconstructed response within a coherent imaging framework.

## Data and Code Availability

Selected processed data and analysis scripts supporting the reported results may be made available by the corresponding authors upon reasonable request. The scope of access and terms of use will be agreed on a case-by-case basis.

## Supporting information

Supplementary Material: Operating Ranges

## A Appendix

### Modal Reference and Radial Phase

#### I. Angular Reference and Modal Blocks

This appendix justifies the two steps used in Section III: Fourier separation of the periodic reference and the high-frequency phase of its radial blocks. With the temporal convention in Section II, outward propagation has positive phase and migration reverses it.

Let Θ = Θ_ap_ and *µ*_*n*_ = 2*πn/*Θ. The object’s Fourier coefficients on *I*_per_ are

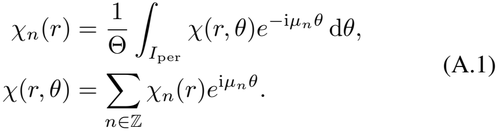

Write *h*(*k*; *r, δ*) for the periodic angular kernel of the reference propagation, including its radial transmit factor. For centered transmission this defines

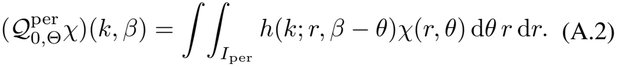

The radial propagation chosen for this reference is derived below. Insert (A.1) and substitute *δ* = *β* − *θ*. Periodicity permits integration over the same interval and gives

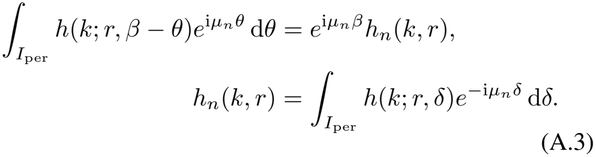

Applying angular analysis to the output, with the 1*/*Θ nor-malization in (A.1), gives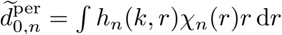. Denote this radial mapby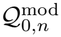. For the unitary transform 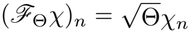,

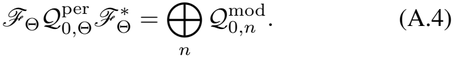

The direct sum expresses independent radial blocks of the periodic reference. The factor 1*/*Θ in angular analysis corresponds to *ω*_*q*_ = 1*/N*_rx_ for uniform samples without apodization in (8). The physical finite-arc model remains (4); the reference supplies the phase used to construct its migration image.

#### II. Radial Equation and Principal Phase

The polar radial equation (9) underlies angular-spectrum propagation [1]. Its outgoing solution is proportional to

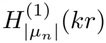. Write its leading oscillatory form as 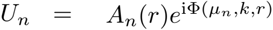. Separating the leading phase and transport terms gives

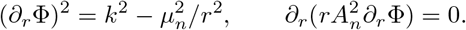

The positive square root specifies outward propagation; the second relation gives the slowly varying amplitude. Integration of the first gives (10). In particular, with *a* = |*µ*_*n*_| and 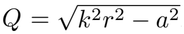

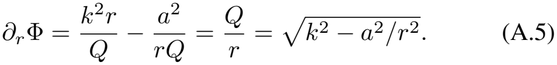

The second term comes from differentiating the inverse cosine. On a branch separated from the turning point, the outgoing radial transfer between *r*_0_ and *r* has principal phase Φ(*µ*_*n*_, *k, r*) − Φ(*µ*_*n*_, *k, r*_0_). Multiplication by 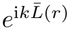 produces (11). The phase-form reference uses

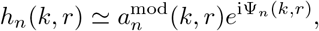

and reconstructs the reference kernel as 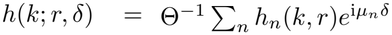. Substitution into the radial block above gives (12). Thus periodic angular separation and the radial high-frequency approximation enter at distinct steps. The image operator reverses this selected reference phase with the fixed migration filter in (15).

## B Appendix

### Phase-Screen Factors and Image Formula

Fix *t, j* as in Section III-C. On the retained modal–spectral support, let 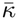 be the midpoint and *H* the half-width of *k*_*n*_(*p*) − *ηp*. Define the normalized spectral residual and the centered path:

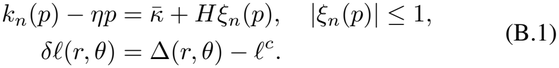

If *H* = 0, set *ξ*_*n*_ = 0.

For real *ζ* = *Hδℓ*, the Jacobi–Anger identity [35] gives

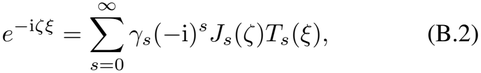

where *J*_*s*_ is the Bessel function, *T*_*s*_(cos *γ*) = cos(*sγ*) is the Chebyshev polynomial, and *γ*_0_ = 1, *γ*_*s*_ = 2 for *s* ≥ 1. Substituting (B.1) into the left-hand side of (22) yields the explicit coefficients

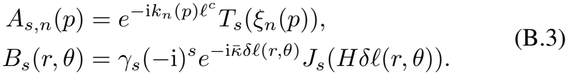

Thus the path-center factor remains wholly in the spectral factor *A*_*s,n*_, and the residual center supplies an exact spatial carrier in the screen *B*_*s*_. Only 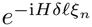 is expanded. The affine approximation assessed in Fig. 4(c) replaces this last factor by one, retaining 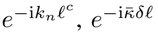, and the query *ρ*. Insert this series into the coupled integral (18). Each *B*_*s*_(*r, θ*) is independent of *n, p* and can be taken outside the modal sum and spectral integral. The remaining integral is precisely *V*_*s*_ in (23), so the truncated local field is 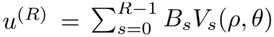 Restoring *t, j* and applying 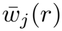 gives (24). For the finite retained mode set and integrable *F*_*n*_ on compact spectral supports, the uniform factorial tail bound on bounded | *Hδℓ*| justifies termwise integration and convergence to the coupled target. Supplementary Section S3.1 supplies that bound.

The shift–residual example in (19) illustrates why this series is needed. At zero offset, 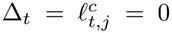, the coefficients satisfy *A*_0,*t,n,j*_ = *B*_0,*t,j*_ = 1 and *B*_*s,t,j*_ = 0 for *s* ≥1, recovering the reference synthesis under the same support and weights. When *H* = 0 with nonzero offset, one term is also sufficient, with the spectral and spatial carriers retained.

