## Supplementary Material: Operating Ranges for "Circular-Arc Modal Migration for Ultrasound Imaging—Part I: Finite-Arc Spectral Theory and Stolt–Pullback Factorization"

This supplement collects implementation guidance, supporting derivations, and numerical checks. Symbols are defined where used; principal notation is listed in Part I, Table I.

#### S1 Implementation Choice and a Worked Example

Table S1 gives starting points from the same-RF comparisons in Section S6. The compact semicircular and conventional convex configurations both image outward from a circular arc. Radii are measured from the curvature center; radial depth is $r - r_0$.

Table S1: Engineering reference for the initial implementation. Bands and radii describe the tested configurations.

| Configuration | Acquisition and image scales | Suggested first trial |
|---|---|---|
| Semicircular array | $r_0 = 0.75$ mm, $r_d = 0.375$ mm; 8–24 MHz; target radii 2.5–5.5 mm; $r_d/r \simeq 0.07$–$0.15$. | Scalar pullback: widths differ from phase screens by less than 5% at the tested targets. |
| Convex arc | $r_0 = 55$ mm, $r_d = 27.5$ mm; about 2.37–4.64 MHz; image radii 60–130 mm; $r_d/r \simeq 0.21$–$0.46$. | Phase screens for shallow and off-axis targets when surrounding-response control is important. |
| Other arrays | Compare relative source displacement, retained band, image radii, and angular support. | Compare both implementations on representative shallow off-axis and deeper data. |

The ratio $|\mu_n|/(kr)$, with $\mu_n = 2\pi n/\Theta_{\mathrm{ap}}$, changes the radial phase slope; source displacement sets the accumulated offset phase. Smaller $r_d/r$ favors scalar pullback for comparable bands and angular support. Compare matched RF, conditioning, transmit weights, and grids against the application's response and runtime requirements.

##### S1.1 Worked Semicircular-Array Accuracy Check

The nominal semicircular example uses 64 receivers, 24 source angles over ±90°, 61 candidate modes, and 713 bins over 8–24 MHz. Its radial plan uses 260 nodes over 1.5–7 mm, a 16–24-MHz design band, and a 0.6-rad local tolerance. Receive/modal weights are rectangular and transmit weights trapezoidal. Screening uses $E^{\mathrm{ker}}_{\mathrm{tot}}(\mathbf{x}) \le \varepsilon_{\mathrm{ker}}$, with the coefficients in Section S4.2. The $66 \times 101$ nominal audit inserts eight radial samples per threshold bracket. All-angle sampled intervals ending at 7 mm start at 5.058–5.068, 3.388–3.397, and 2.519–2.529 mm for tolerances 0.1, 0.2, and 0.3. These brackets describe that grid.

Matched modal point tests clarify what the coefficient check means for a response. With $\phi_{tn}(k;\mathbf{x}) = \Psi^0_n(k,r) + k\Delta_t(\mathbf{x})$, the direct reference is

$$h_{\mathrm{dir}}(\mathbf{x};\mathbf{x}_*) = \sum_{t,n,i,j} \omega^{\mathrm{ker}}_{tnji}(\mathbf{x}) e^{-\mathrm{i}[\phi_{tn}(k_i;\mathbf{x}) - \phi_{tn}(k_i;\mathbf{x}_*)]} e^{\mathrm{i}2n(\theta-\theta_*)}. \tag{S1}$$

The scalar phase proxy substitutes the local phase of Section S2.2; the interpolated scalar evaluation also performs Stolt interpolation and radial synthesis, removing and restoring a 20-MHz carrier around interpolation. All three share modal point inputs and amplitude marginals, distinct from channel-RF tests.

At twenty targets, cuts span ±0.45 mm radially and ±1.2 mm tangentially, sampled at 0.005 and 0.01 mm, without peak alignment. Relative to the direct reference, the largest scalar displacement is 0.051 reference FWHM, width change 11.83%, and peak-amplitude change 1.005 dB. All twenty meet displacement ≤ 0.25 FWHM and width change ≤ 15%; fifteen also meet amplitude change ≤ 1 dB and outer-response change ≤ 3 dB. Outer response is the peak-normalized maximum beyond one reference FWHM within each cut. The interpolated response adds radial broadening and attenuation to the phase proxy; halving cut spacing changes FWHM by at most 0.32%. Thus kernel and response checks supply complementary information.

### S2 Reference Mapping and the Scalar Implementation

For reference slope $\lambda_j = \bar{L}'(r_j) > 0$ and $b_{n,j} = \mu_n^2/r_j^2$, the map $p = \lambda_j k + \sqrt{k^2 - b_{n,j}}$ is strictly increasing on $k > \sqrt{b_{n,j}}$. Choosing the quadratic root with $p - \lambda_j k > 0$ gives

$$k_{n,j}(p) = \frac{p^2 + b_{n,j}}{\sqrt{p^2 + (1-\lambda_j^2)b_{n,j}} + \lambda_j p}, \qquad D_{n,j} = \left(\lambda_j + \frac{k_{n,j}}{\sqrt{k_{n,j}^2 - b_{n,j}}}\right)^{-1}. \tag{S2}$$

The image integral changes measure by $\mathrm{d}k = D_{n,j}\mathrm{d}p$, on $p > \lambda_j\sqrt{b_{n,j}}$. At $\lambda_j = 1$ this recovers the main-paper Stolt inverse.

#### S2.1 Semicircular Reference and Geometrical Fit

Set $\vartheta_t = \theta - \alpha_t$, $\bar{L} = r$, and $\Delta_t = L_{\mathrm{tx},t} - r$. The second-order offset path separates as

$$\Delta_t^{(2)} = \frac{r_d^2}{4r} - r_d\cos\vartheta_t - \frac{r_d^2}{4r}\cos 2\vartheta_t, \quad \lambda_c(r) = 1 - \frac{r_d^2}{4r^2}, \quad p_{n,j}^{\mathrm{sc}} = \lambda_c(r_j)k + \sqrt{k^2 - (2n)^2/r_j^2}. \tag{S3}$$

The phase anchor remains the centered-source value $\Psi_n^0(k, r_j) = kr_j + \Phi(2n, k, r_j) - \Phi(2n, k, r_0)$. Thus its local phase uses $\Psi_n^0(k, r_j) + p_{n,j}^{\mathrm{sc}}(r - r_j)$, retaining the implemented anchor–slope convention. Let $g_q = (r - r_0\cos(\theta - \beta_q))/L_{\mathrm{rx},q}$ and $\lambda_q^{\mathrm{path}} = \lambda_c + g_q$. Minimizing $\sum_q \langle (1 - \lambda\lambda_q^{\mathrm{path}})^2\rangle_\theta + \varepsilon_{\mathrm{fit}}\lambda^2$ gives

$$\lambda_{\mathrm{sc}}(r) = \frac{\sum_q \langle \lambda_q^{\mathrm{path}}\rangle_\theta}{\sum_q \langle (\lambda_q^{\mathrm{path}})^2\rangle_\theta + \varepsilon_{\mathrm{fit}}}, \qquad \rho_t^{\mathrm{sc}} = r + \lambda_{\mathrm{sc}}(r)\Delta_t^{(2)}. \tag{S4}$$

The semicircular array uses equal receiver weights, a uniform image-angle mean, and $\varepsilon_{\mathrm{fit}} = 10^{-12}$. This fit is geometrical and independent of the RF. With the weighted Stolt spectrum $F_{t,n,j}^{\mathrm{sc}}$ of Part I and mode-dependent overlap $w_{n,j}^{\mathrm{sc}}$, the reference image is

$$V_t^{\mathrm{sc}}(\rho, \theta) = \sum_n e^{\mathrm{i}2n\theta} \frac{\sum_j w_{n,j}^{\mathrm{sc}}(\rho) \int_{P_{n,j}^{\mathrm{sc}}} F_{t,n,j}^{\mathrm{sc}}(p) e^{-\mathrm{i}p\rho}\,\mathrm{d}p}{\sum_j w_{n,j}^{\mathrm{sc}}(\rho)}. \tag{S5}$$

Zero-denominator fields vanish. Overlap is evaluated at the query radius before angular summation; the prepared domain covers all queries.

#### S2.2 Signed Residual and Scalar Exactness

For $\rho = \rho_t^{\mathrm{sc}}$, subtract the exact-path modal phase from the implemented phase:

$$\begin{aligned} e_{\mathrm{loc}} &= \Psi_n^0(k, r_j) + p_{n,j}^{\mathrm{sc}}(\rho - r_j) - \Psi_n^0(k, \rho), & e_{\mathrm{TX}} &= k(\Delta_t^{(2)} - \Delta_t), \\ e_{\mathrm{pb}} &= \Psi_n^0(k, \rho) - \Psi_n^0(k, r) - k\Delta_t^{(2)}, & e_{\mathrm{tot}} &= e_{\mathrm{loc}} + e_{\mathrm{TX}} + e_{\mathrm{pb}}. \end{aligned} \tag{S6}$$

Here loc, TX, and pb identify local radial synthesis, transmit-path expansion, and scalar pullback. Writing $p_{n,j}^0 = \partial_r \Psi_n^0(k, r_j)$ gives

$$e_{\mathrm{loc}} = \underbrace{\Psi_n^0(k, r_j) + p_{n,j}^0(\rho - r_j) - \Psi_n^0(k, \rho)}_{\text{quadratic Taylor remainder}} + k[\lambda_c(r_j) - 1](\rho - r_j). \tag{S7}$$

The second term is linear in query-to-center distance and is retained in the scalar audit.

For a nonzero fixed $\Delta_t$ and frequency-independent carrier $a$ and shift $s$, exact equality $e^{-\mathrm{i}k_n(p)\Delta_t} = ae^{-\mathrm{i}ps}$ for every spectrum on an open $p$ interval requires $k_n'(p)\Delta_t = s$, by differentiation. Thus $k_n$ must be affine; a shared query requires a common slope, and one common carrier requires coincident carrier factors. For unit reference slope, $k_n''(p) = b_{n,j}/p^3$. The geometrical fit above approximates the offset phase through its fitted query; the phase-screen expansion controls the residual after its own query construction.

### S3 Phase-Screen Bounds and the Planar Limit

On the compact retained support $\mathcal{S}_j = \{(n, p) : p \in P_{n,j}\}$, choose

$$\eta_j \in \arg\min_{\eta}\{\max_{\mathcal{S}_j}(k_{n,j} - \eta p) - \min_{\mathcal{S}_j}(k_{n,j} - \eta p)\}. \tag{S8}$$

This objective is convex. Two distinct $p$ values give the coercive lower bound $|\Delta k - \eta\Delta p|$; if all $p$ coincide, the width is constant. The extrema define midpoint $\bar{\kappa}_j$ and half-width $H_j$, so $k_{n,j} = \eta_j p + \bar{\kappa}_j + H_j\xi_{n,j}$ with $|\xi_{n,j}| \leq 1$. For synthesis domain $[a_j, b_j]$, choose the residual-path center

$$\ell^c_{t,j} = \operatorname{proj}_{\mathcal{C}_{t,j}} \frac{\Delta_{\max} + \Delta_{\min}}{2}, \qquad \mathcal{C}_{t,j} = \bigcap_{(r,\theta)\in I_j\times I_\theta} \{c : a_j \leq r + \eta_j(\Delta_t - c) \leq b_j\}. \tag{S9}$$

The midpoint minimizes maximum path deviation, and projection gives the constrained minimum. An empty feasible interval requires a larger synthesis domain. With $\delta\ell = \Delta_t - \ell^c$ and $\rho = r + \eta_j\delta\ell$,

$$e^{-\mathrm{i}pr - \mathrm{i}k_{n,j}\Delta_t} = e^{-\mathrm{i}k_{n,j}\ell^c} e^{-\mathrm{i}\bar{\kappa}_j\delta\ell} e^{-\mathrm{i}H_j\delta\ell\xi_{n,j}} e^{-\mathrm{i}p\rho}.$$

#### S3.1 Truncation and Radial Error

Only the third factor is truncated. For $|\zeta| = |H_j\delta\ell| \leq Z$ and $R \geq 1$, the Jacobi–Anger/Chebyshev series of Part I, Appendix B, and $|T_s(\xi)| \leq 1$ give

$$\epsilon_R(\zeta) \leq 2\sum_{s=R}^{\infty} |J_s(\zeta)| \leq 2\sum_{s=R}^{\infty} \frac{(Z/2)^s}{s!} \leq \frac{2(Z/2)^R}{R!\,[1 - Z/(2(R+1))]}. \tag{S10}$$

The last inequality additionally requires $R + 1 > Z/2$. The bound $|J_s(\zeta)| \leq (|\zeta|/2)^s/s!$ follows by taking absolute values in the Poisson integral for $J_s$ and evaluating its beta integral. The factorial majorant gives uniform convergence on bounded $\zeta$ and $|\xi| \leq 1$, justifying termwise integration for integrable $F_{t,n,j}$. A uniform kernel error $\epsilon$ yields

$$|u_{t,j} - u^{(R)}_{t,j}| \leq \epsilon \sum_n \int_{P_{n,j}} |F_{t,n,j}(p)|\,\mathrm{d}p. \tag{S11}$$

For positive quadrature, replace the integral by its weighted sum and enclose the retained nodes. The other factors have unit modulus. The reference phase has second derivative $\partial_r^2\Psi_n = k\bar{L}''(r) + \mu_n^2/[r^3\sqrt{k^2 - \mu_n^2/r^2}]$. On an interval of half-width $h_j$ separated from the turning point,

$$|\Psi_n(k, r) - \Psi_n(k, r_j) - \partial_r\Psi_n(k, r_j)(r - r_j)| \leq \tfrac{1}{2}h_j^2 \sup_{\varrho\in I_j} |\partial_r^2\Psi_n(k, \varrho)|. \tag{S12}$$

The scalar implementation additionally retains the linear anchor term in (S7).

#### S3.2 Local Planar Limit

Let $r_0 = R \to \infty$ with physical aperture length $L$ fixed, $r = R + z$, $\theta = x/R$, and $\mu_n = 2\pi nR/L$. Then $\kappa_x = \mu_n/R$ stays finite and $e^{\mathrm{i}\mu_n\theta} = e^{\mathrm{i}\kappa_x x}$. For bounded $z$ and spectral support uniformly separated from $k = |\kappa_x|$,

$$\Phi(\mu_n, k, R + z) - \Phi(\mu_n, k, R) = \int_0^z \sqrt{k^2 - \frac{\kappa_x^2 R^2}{(R + s)^2}}\,\mathrm{d}s \longrightarrow z\sqrt{k^2 - \kappa_x^2}. \tag{S13}$$

With unit transmit slope, $p \to k + \sqrt{k^2 - \kappa_x^2}$ and $k = (p^2 + \kappa_x^2)/(2p)$. A virtual source fixed at $(x_s, -D)$ requires $r_d = R - D$ and $\alpha = x_s/R$; its residual from $\bar{L} = r - r_d$ tends to $\sqrt{(x - x_s)^2 + (z + D)^2} - (z + D)$. The planar limit thus recovers Cartesian propagation while leaving a finite-distance offset correction. Amplitudes, aperture support, and interpolation remain separate operator choices.

### S4 Implemented Weights and Error-Audit Conventions

#### S4.1 Weights and Semicircular Phase Anchor

The density $D_{n,j}$ changes measure, $W_{n,j}$ restricts support, and normalized square-root-Hann windows blend local fields. Migration also weights by

$$a_{n,j}^{\mathrm{im}}(p) = \frac{D_{n,j}(p)}{D_{n,j}(p)^2 + \lambda_{\mathrm{amp}}[1 + (p/p_{*,n,j})^2]}, \quad \lambda_{\mathrm{amp}} = 10^{-3}. \tag{S14}$$

Here $p_{*,n,j} = 1$ rad/m for the semicircular array and median local $|p|$ for the convex arc. Baselines use unit modal and receive tapers. The semicircular taper control uses $\mathrm{sinc}(n/31)$ for $|n| \leq 30$, where $\mathrm{sinc}(x) = \sin(\pi x)/(\pi x)$; convex C1 uses 150 unit and 42 endpoint-taper receive coefficients for both methods. The unit post-Stolt control sets $a_{n,j}^{\mathrm{im}} = 1$. Normalized trapezoidal transmit weights are $v_1 = v_{24} = 1/46$ and $v_t = 1/23$ otherwise. Both methods share a convex RF cosine taper covering 10% of band samples in total.

Semicircular support requires $kr_j \geq \pi$ and $k^2 r_j^2 > 3\mu_n^2$. For nonzero mode and $\varepsilon = 10^{-12}$, the implemented receive anchor is

$$\Phi_{\mathrm{clip}}(\mu_n, k, r_0) = \sqrt{\max(k^2 r_0^2 - \mu_n^2, 0)} - |\mu_n| \cos^{-1}(\min\{1 - \varepsilon, |\mu_n|/(kr_0 + \varepsilon)\}) \,. \tag{S15}$$

The zero-mode value is $kr_0$. The anchor uses a guarded propagating phase within that branch and a numerical continuation outside it. Its independence of image radius preserves the local Stolt derivative. A modal phase multiplier corrects the angular origin in both images and audits.

#### S4.2 Discrete Scalar-Audit Coefficients

For radial interpolation nodes $r_-, r_+$ around $\rho$, with linear interpolation weights $\omega_\nu^{\mathrm{int}}$, define

$$\begin{aligned} \omega_{tnj}^{\mathrm{rad}}(\mathbf{x}) &= \sum_{\nu \in \{-,+\}} \omega_\nu^{\mathrm{int}}(\rho) \frac{w_{n,j}^{\mathrm{sc}}(r_\nu)}{\sum_{j'} w_{n,j'}^{\mathrm{sc}}(r_\nu)}, \\ \omega_{tnji}^{\mathrm{ker}} &= v_t \omega_n^{\mathrm{ang}} \omega_{tnj}^{\mathrm{rad}} D_{n,j}(p_{n,j}^{\mathrm{sc}}(k_i)) \sum_l \Pi_{nj,li}^{\mathrm{St}} a_{n,j}^{\mathrm{im}}(p_l) \frac{\Delta p_l}{2\pi}. \end{aligned} \tag{S16}$$

Zero-denominator contributions vanish. Here $\Pi_{nj,li}^{\mathrm{St}}$ maps native frequency $k_i$ to Stolt node $p_l$, $\Delta p_l > 0$ is its quadrature weight, and $\omega_n^{\mathrm{ang}} = 1$ in the maps. These nonnegative amplitude marginals collect interpolation and quadrature; the audit replaces their oscillatory factors by the analytical phase at $\rho$.

Each signed residual in (S6) forms a vector over $(t, n, i)$ after summing active intervals:

$$\begin{aligned} [\mathbf{h}_{\mathrm{test}}]_{tni} &= \sum_j \omega_{tnji}^{\mathrm{ker}} e^{-\mathrm{i} e_{\mathrm{test}}}, \qquad [\mathbf{h}_{\mathrm{ref}}]_{tni} = \sum_j \omega_{tnji}^{\mathrm{ker}}, \\ (E_{\mathrm{test}}^{\mathrm{ker}})^2 &= \frac{\|\mathbf{h}_{\mathrm{test}}\|_2^2 + \|\mathbf{h}_{\mathrm{ref}}\|_2^2 - 2|\mathbf{h}_{\mathrm{ref}}^* \mathbf{h}_{\mathrm{test}}|}{\|\mathbf{h}_{\mathrm{ref}}\|_2^2}. \end{aligned} \tag{S17}$$

This discrepancy removes one common phase from the compounded coefficient row, preserving relative phases across modes, frequencies, events, and intervals, including interval cancellation.

#### S4.3 Phase-Screen Maps

Let $\mathcal{J}(r) = \{j : w_j(r) > 0\}$ and $\zeta_{t,j} = H_j(\Delta_t - \ell_{t,j}^c)$. The four main-paper phase-screen maps are

$$\begin{aligned} E_{\mathrm{aff}} &= \max_{t,j \in \mathcal{J}(r)} |\zeta_{t,j}|, \qquad E_{\mathrm{aff}}^{\mathrm{ker}} = \max_{t,j \in \mathcal{J}(r)} 2 \sin\left(\frac{\min\{|\zeta_{t,j}|, \pi\}}{2}\right), \\ E_{\mathrm{screen}} &= \max_{t,j \in \mathcal{J}(r)} 2 \sum_{s=R_{t,j}}^{\infty} |J_s(\zeta_{t,j})|, \quad R_{\max} = \max_{t,j \in \mathcal{J}(r)} R_{t,j}. \end{aligned} \tag{S18}$$

The affine kernel envelope equals $\sup_{|\xi| \leq 1} |e^{-\mathrm{i}\zeta\xi} - 1|$ and saturates at two for $|\zeta| \geq \pi$. The screen error is bounded over the continuous residual interval enclosing the retained Stolt nodes. $R_{t,j}$ counts terms $s = 0, \ldots, R_{t,j} - 1$.

### S5 Numerical and Weighting Controls

The controls isolate approximation and evaluation steps using the support and conventions above. For the semicircular array, query-centered interval refinement makes $e_{\mathrm{loc}} \to 0$ while $e_{\mathrm{tot}} \to \Psi_n^0(k, \rho_t^{\mathrm{sc}}) - \Psi_n^0(k, r) - k\Delta_t$. The linear anchor term in (S7) therefore changes the refinement order, whereas the remaining scalar/path error persists. Table S2 summarizes these checks and the convex direct-integral verification.

Table S2: Numerical checks on the reported reconstruction plans. Relative field errors compare complex values without gain or phase alignment.

| Check | Observed result and comparison target |
|---|---|
| Semicircular radial refinement | Half-width 0.1 to 0.0015625 mm reduces the implemented local phase residual from $2.313 \times 10^{-2}$ to $3.729 \times 10^{-4}$ rad. Its leading term is linear; a consistently centered Taylor control converges quadratically. |
| Transmit expansion | Source-radius refinement gives remainder orders 3.053–3.002, consistent with a second-order path expansion. |
| Convex radial bound | The maximum residual over 19 intervals is 0.2722 rad for the 0.3-rad criterion. |
| Convex screen truncation | Planned 4–17 terms give maximum continuous tail bound $1.243 \times 10^{-3}$ and maximum sampled factor error $9.914 \times 10^{-4}$. |
| Convex direct integral | Across four $3 \times 3$ coordinate blocks, the separated versus direct local integral has relative errors $2.404 \times 10^{-5}$–$1.985 \times 10^{-4}$ with planned terms; at 24 terms the maximum is $2.525 \times 10^{-10}$. |
| Semicircular screen evaluation | Increasing 10 to 12 terms changes the full image by $9.73 \times 10^{-10}$ in relative norm. Direct quadrature at 60 target/background pixels agrees to $1.23 \times 10^{-7}$. |

The convex direct check uses the same locally linearized integral, inverse maps, amplitudes, overlap, C1 receive taper, and coherent compounding in both calculations: 24 transmissions, 638 frequency bins, 191 candidate modes, and 256 common radial-wavenumber nodes per interval. Radial Fourier sums are evaluated directly at queries. Block centers $(r - r_0, \theta)$ are approximately $(6.92\,\mathrm{mm}, -24.92^\circ)$, $(9.94, 0^\circ)$, $(31.90, -12.03^\circ)$, and $(70.06, 0^\circ)$. This tests numerical evaluation. The same-RF point-response comparison is summarized separately in Section S6.

#### S5.1 One-Factor Weighting Comparisons

Receive/modal taper and the post-Stolt amplitude are varied separately, keeping RF conditioning, grid, transmit quadrature, and phase operations fixed. Removing the semicircular Lanczos taper narrows tangential FWHM by 13.5–15.7%, with unchanged peak cells. Convex widths change little under matched C1 receive taper. Unit post-Stolt controls leave principal widths similar; semicircular radial profiles change more visibly below −30 dB. The migration weighting and phase operations therefore contribute on different response scales.

Table S3: FWHM ranges for the semicircular (20 targets) and convex (4 targets) configurations, in mm. CAMM denotes the respective scalar and phase-screen baselines.

| Configuration | Radial | Tangential |
|---|---|---|
| Semicircular DAS | 0.113–0.118 | 0.138–0.459 |
| Semicircular CAMM, untapered | 0.117–0.122 | 0.158–0.494 |
| Semicircular CAMM, Lanczos | 0.116–0.120 | 0.186–0.580 |
| Semicircular CAMM, unit post-Stolt | 0.115–0.120 | 0.156–0.489 |
| Convex DAS, rectangular / C1 | 0.587–0.677 / 0.590–0.677 | 0.276–0.687 / 0.278–0.726 |
| Convex CAMM, rectangular / C1 | 0.637–0.682 / 0.637–0.683 | 0.430–0.906 / 0.430–0.907 |
| Convex CAMM, unit post-Stolt | 0.638–0.682 | 0.436–0.909 |

### S6 Same-RF Point-Response Comparisons

Within each geometry, DAS, phase-screen CAMM (PS), and scalar-pullback CAMM (SP) share RF, timing, output grid, retained band, rectangular receive weights, unit modal taper, and trapezoidal transmit weights. CAMM implementations retain their respective radial plans, interpolation, and overlap. Images retain their original coordinates; displays use whole-field peak normalization, with semicircular angular cuts normalized per cut.

Semicircular DAS uses a length-8900 FFT and the CAMM band (98.894% of positive-frequency RF energy). CAMM uses 713 frequencies, 61 modes, and the clipped anchor. SP blends reference fields at the shifted radius with a second-order path; PS uses the exact path and square-root-Hann overlap at the physical radius, assembling mode-dependent intervals before angular summation. PS chirp-z fields use eightfold radial sampling and carrier-demodulated cubic interpolation. Archived timing is retained; excitation/time-origin metadata are unavailable.

The convex routes use $\bar{L} = r - r_d$ and exact $L_{\mathrm{tx},t}$. SP fits $\lambda_{\mathrm{sc}} = \langle a \rangle / (\langle a^2 \rangle + 10^{-12})$, where $a = 1 + \partial_r L_{\mathrm{rx}}$, using normalized receiver and trapezoidal image-angle quadrature. It queries $r + \lambda_{\mathrm{sc}}[L_{\mathrm{tx},t} - (r - r_d)]$. The coefficient spans 0.5081–0.6465; queries span 60.0000–138.7952 mm with all 24 events valid at every output pixel. Synthesis grids are prepared separately.

Semicircular peak searches/cuts span $\pm 0.4$ mm and $\pm 10°$ about each target. Convex searches use $\pm 2.5$ mm and $\pm 3°$, with width cuts of $\pm 4$ mm and $\pm 6°$ about the detected peak. Interpolated half-amplitude crossings define FWHM; tangential width is angular width times detected radius. At the twenty targets in the semicircular configuration, PS widths are 1.47–3.61% smaller radially and 1.09–4.94% smaller tangentially than SP. Radial peak indices coincide; angular indices coincide at fourteen targets and differ by one pixel at six. Both have maximum angular deviation 0.30°. The locations are deterministic, within one acquisition per geometry.

#### S6.1 Matched Shallow Axis/Off-Axis Pair

Independent points at $r - r_0 = 10$ mm and $\theta = 0°, -20°$ share a 55-mm array radius, 27.5-mm source radius, 192 receivers, 24 transmissions, excitation, and 100-MHz sampling (16 000 samples; $-1.24$ $\mu$s pulse-center correction). The off-axis coordinate is $(-22.231, 6.080)$ mm, with apex-referenced $z$. Reconstructions use 2.3625–4.6375 MHz and a $256 \times 257$ grid over $r = 60$–130 mm and $\theta = \pm 27.5°$.

The full-field outside-neighborhood energy fraction sums $|u|^2 r \Delta r \Delta\theta$ outside a fixed neighborhood, divided by the sum over the whole common reconstruction domain. The neighborhood spans $\pm 1$ mm radially and $\pm 3$ mm tangentially about the true target, with tangential distance $r_{\mathrm{target}}(\theta - \theta_{\mathrm{target}})$. Table S4 shows the widths and wider-scale response together. All methods locate each target in the same pixel; PS has lower outside fraction and wider FWHM than DAS.

Table S4: Shallow-pair measurements on the common grid. FWHM and peak error are in mm; outside energy is a percentage of the full-field image energy.

| Angle | Method | Peak error | Radial FWHM | Tangential FWHM | Outside energy |
|---|---|---|---|---|---|
| 0° | DAS | 0.059 | 0.587 | 0.276 | 4.14 |
| | PS | 0.059 | 0.638 | 0.430 | 1.36 |
| | SP | 0.059 | 0.625 | 0.532 | 13.47 |
| −20° | DAS | 0.063 | 0.591 | 0.295 | 3.45 |
| | PS | 0.063 | 0.639 | 0.472 | 0.61 |
| | SP | 0.063 | 0.631 | 0.517 | 11.82 |

#### S6.2 Planning and Online Work

Fixed geometry, band, and grid permit reusable weights and transform plans. Online processing still performs RF analysis, remapping, synthesis, queries, and compounding. Phase screens synthesize $R_{t,j}$ fields per event/interval; scalar pullback queries one reference image. For $P_j$ spectral and $Q_j$ query nodes, radial sums cost $O(P_j Q_j)$ per right-hand side, or $O(r_j^{\mathrm{rad}}(P_j + Q_j))$ with rank-$r_j^{\mathrm{rad}}$ factorization. Part II measures complete implementations on matched hardware.
